# The transcription factor BATF pioneers the differentiation program of cytolytic effector CD8^+^ T cells through the direct interaction with IRF4

**DOI:** 10.1101/2024.08.21.605975

**Authors:** Sotaro Fujisawa, Yamato Tanabe, Arisa Hojo, Ryotaro Shiga, Junko Kurachi, Miki Koura, Toshikatsu Tamai, Yusuke Miyanari, Edward John Wherry, Makoto Kurachi

**Author notes:** Correspondence should be addressed to: Makoto Kurachi. These authors contributed equally (Sotaro Fujisawa, Yamato Tanabe).

## Abstract

The transcription factor BATF plays critical roles in the differentiation of various immune cells, including CD8^+^ T cells. Here, we demonstrated that BATF controls epigenomic and transcriptomic reprogramming of CD8^+^ T cells at an early phase of acute viral infection, thereby promoting the differentiation of cytolytic effector CD8^+^ T cells. Loss of BATF drastically perturbed gene expression, chromatin accessibility, and the bindings of key transcription factors including Jun, T-bet, and IRF4. The direct interaction with IRF4 was essential for BATF-mediated effector differentiation, as the BATF mutant lacking this interaction failed to induce proper chromatin remodeling and proliferation of antigen-specific CD8^+^ T cells. Notably, IRF4 binding was exhaustively dependent on BATF, whereas BATF retained binding capacity even in IRF4-deficient CD8^+^ T cells. Furthermore, BATF initiated chromatin remodeling in the absence of IRF4, whereas subsequent dynamic epigenomic reorganization required IRF4. Our data proposed that BATF serves as a “pioneer transcription factor” spearheading the reorganization of chromatin architecture upon antigen encounter, followed by further rearrangement of epigenomic and transcriptomic landscapes through the cooperation with IRF4.

## Introduction

Upon activation by their cognate antigens, naive CD8^+^ T cells undergo clonal, exponential proliferation, and differentiate into effector CD8^+^ T cells. During the earliest phase of effector differentiation, CD8^+^ T cells initially differentiate into early effector cells (EEC) characterized by low expression of both killer lectin-like receptor KLRG1 and interleukin 7 receptor alpha (CD127), followed by further differentiation into two distinct subsets, short-lived effector cells (SLEC; KLRG1^hi^CD127^lo^) and memory precursor cells (MPEC; KLRG1^lo^CD127^hi^) (Joshi et al., 2007; Kaech et al., 2003; Kaech & Cui, 2012; Obar et al., 2011). SLEC exhibit greater proliferative and cytotoxic capacities and play a pivotal role in eliminating infected cells. After pathogen clearance, a vast majority of SLEC undergoes apoptosis, whereas a small fraction persists and differentiates into effector-memory or tissue-resident memory populations (Herndler-Brandstetter et al., 2018). In contrast, MPEC are less proliferative and cytotoxic but harbor greater potential to persist and form a pool of memory cells (Huster et al., 2004; Kaech et al., 2003; Yuzefpolskiy et al., 2015). These fate decisions during early phase of infection are governed by the strength of signals from the T cell receptor (TCR), co-stimulation, and cytokine-mediated inflammation, followed by transcriptional changes regulated by transcription factors (TFs) (Kaech & Cui, 2012). For instance, T-box transcription factor 21 (T-bet), inhibitor of binding 2 (Id2), and B lymphocyte-induced maturation protein 1 (Blimp-1) promote the differentiation of SLEC, whereas Id3, eomesdermin (Eomes), T cell factor 1 (TCF-1), and forkhead box O1 (FoxO1) control MPEC differentiation (Jeannet et al., 2010; Joshi et al., 2007; Kim et al., 2013; Rutishauser et al., 2009; Yang et al., 2011; Zhou et al., 2010).

Basic leucine zipper transcription factor ATF-like (BATF) is a member of activator protein 1 (AP-1) family transcription factor with a crucial role in regulating differentiation and function of hematopoietic lineages, including lymphocytes. In CD4^+^ T cells, BATF regulates the differentiation of interleukin 17-producing helper T cells (TH17 cells) (Ciofani et al., 2012; Pham et al., 2023; Schraml et al., 2009), and FoxP3^+^ regulatory T cells (Hayatsu et al., 2017; Itahashi et al., 2022). BATF is also essential for the development of follicular helper T cells (TFH cells) and for class-switch recombination in B cells (Ise et al., 2011). Importantly, BATF initiates TH17 cell differentiation through interaction with interferon regulatory factor 4 (IRF4) (Ciofani et al., 2012; Murphy et al., 2013). In CD8^+^ T cells, BATF is indispensable during the early stages of effector differentiation, with BATF-deficient CD8^+^ T cells being defective in proliferation and displaying aberrant expression of cytokines and other TFs (Kurachi et al., 2014). A recent study also demonstrated that BATF organizes the bindings of other TFs in CD8^+^ T cells activated *in vitro* (Tsao et al., 2022), suggesting that BATF plays a central role in transcriptional and epigenomic reprogramming in activated CD8^+^ T cells. Moreover, in *in vitro*-activated CD8^+^ T cells, a large portion of BATF-bound genomic regions was co-localized by IRF4, suggesting that the cooperation between BATF and IRF4 is critical also in CD8^+^ T cells (Kurachi et al., 2014; Tsao et al., 2022). Nevertheless, the molecular mechanisms by which BATF regulates epigenomic programs and how BATF and IRF4 (and other TFs) cooperatively function during early phase of acute viral infection remain incompletely understood.

In the present study, we investigated the role of BATF in initiating CD8^+^ T cell effector differentiation *in vivo* using mice models of acute viral infection. We found that BATF was critical for the differentiation and proliferation of cytolytic effector CD8^+^ T cells. Specifically, BATF was required for the rearrangement of chromatin architectures in response to antigens and subsequent epigenomic and transcriptional modification, achieved through orchestrating the bindings of several key TFs. Particularly, we found that the interaction between BATF and IRF4 was critical to initiate chromatin remodeling, and furthermore, the binding of IRF4 necessitated a direct interaction with BATF, while BATF bound to DNA in an IRF4-independent manner. Overall, our findings demonstrate that BATF “pioneers” the effector differentiation of CD8^+^ T cells.

## Results

### BATF is a critical regulator of cytolytic effector CD8^+^ T cell differentiation

To better understand how the loss of BATF impacts the phenotype of CD8^+^ T cells during the early stages of effector differentiation, we adoptively transferred congenically distinct WT and *Batf*^-/-^ P14 CD8^+^ T cells (TCR transgenic that are specific for the lymphocytic choriomeningitis virus (LCMV) gp33 epitope) at a ratio of 1:1 into naive WT recipients, followed by intraperitoneal infection of host mice with Armstrong (Arm) strain of LCMV (Supplementary figure 1A). As reported previously (Kurachi et al., 2014), clonal expansion of *Batf*^-/-^ P14 cells was significantly impaired at day 5 post-infection (Figs. 1A and B). We also analyzed the phenotypes of each P14 cell and observed that approximately 80% of WT P14 cells differentiated into Tim-3^hi^CD62L^lo^ cytolytic effector cell subset, while the proportion of Tim-3^hi^CD62L^lo^ cells remained around 50% in *Batf*^-/-^ P14 cells (Fig. 1C). In contrast, the Tim-3^lo^CD62L^hi^ subset, which corresponds to naive or memory precursor subsets, was more frequently found in *Batf*^-/-^ P14 cells. Meanwhile, the number of Tim-3^lo^CD62L^hi^ population was comparable between WT and *Batf*^-/-^ P14 cells, indicating that differentiation of this subset was not affected by the loss of BATF (Fig. 1D). We also noted reduced expression of several surface proteins (PD-1, CD122, CD49d) and nuclear proteins (T-bet, IRF4, Ezh2) representing cytolytic effector cells in *Batf*^-/-^ P14 cells, whereas the expression of TCF-1, the transcription factor required for the differentiation and maintenance of MPEC (J. Zhang et al., 2021), was significantly higher in *Batf*^-/-^ P14 cells (Fig. 1E and supplementary figures 1B, C). Furthermore, Tim-3 expression was significantly higher in BATF^hi^ WT P14 cells compared to that in BATF^lo^ P14 cells (Fig. 1F, Supplementary figure 1D). Collectively, these data indicate that BATF is a critical for the differentiation of cytolytic effector CD8^+^ T cells during acute viral infection.

**Fig. 1.**
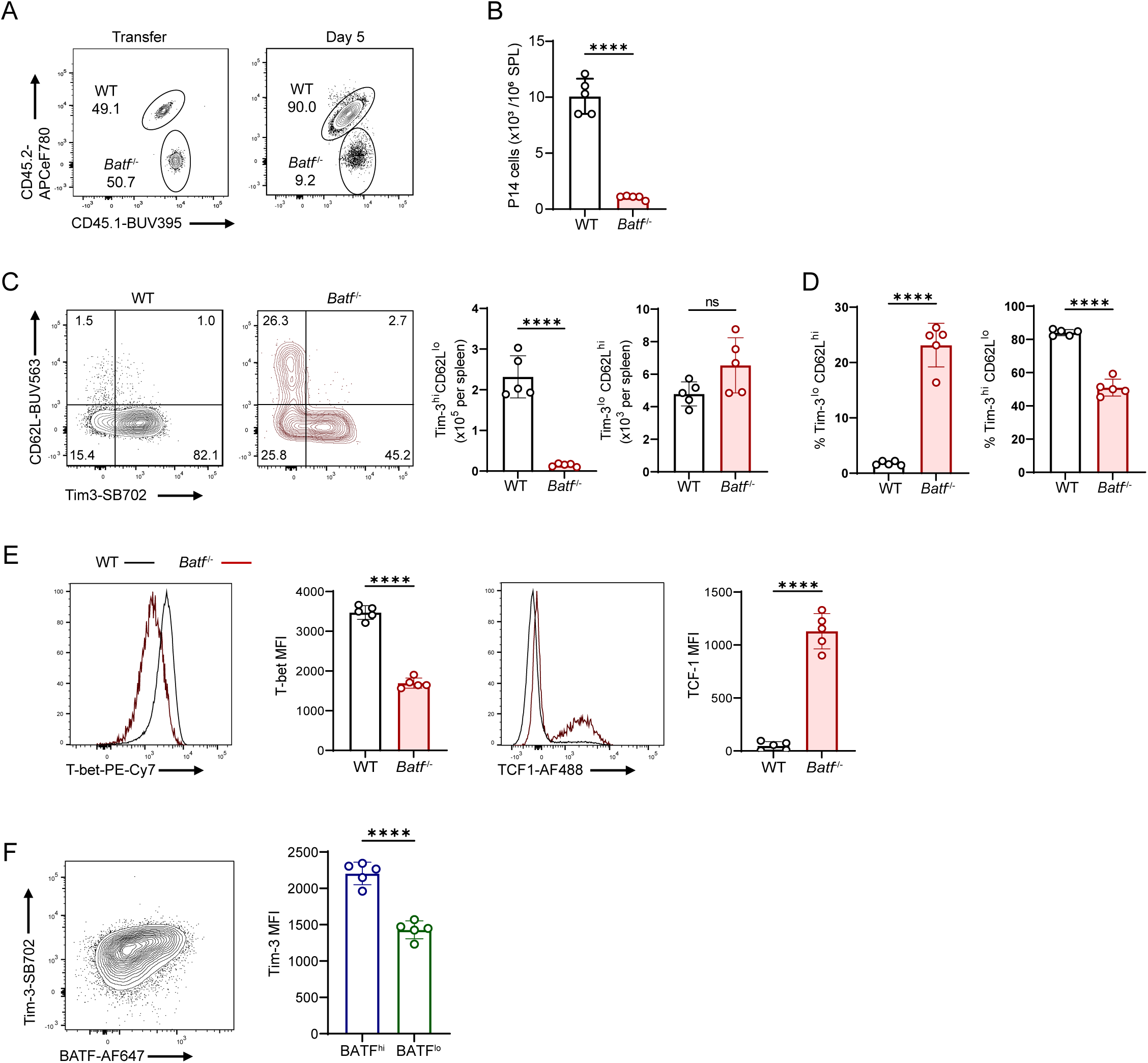
BATF is required for the differentiation of cytolytic effector CD8^+^ T cell during acute viral infection. (A) Flow cytometric analysis of cells obtained from spleen of WT recipient mice adoptively transferred with congenically distinct WT and *Batf*^-/-^ P14 cells mixed at 1:1 ratio (1 × 10^4^ each cell), followed by infection of the recipient mice with LCMV Arm, analyzed at transfer (left) and day 5 post-infection (right plot). Plots are gated on P14 cells. (B) The numbers of WT and *Batf*^-/-^ P14 cells in the spleen of recipient mice in A. (C) Expression of Tim-3 and CD62L at day 5 post-infection in the recipient mice in A. Bar plots indicate the frequencies of Tim-3^hi^CD62L^lo^ (left) and Tim-3^lo^CD62L^hi^ (right) populations in each P14 cell. (D) Quantification of Tim-3^hi^CD62L^lo^ (left) and Tim-3^lo^CD62L^hi^ (right) P14 cells in the spleen of recipient mice in A. (E) Expression of T-bet and TCF-1 at day 5 post-infection in the recipient mice in A. Bar plots represent mean fluorescence intensity (MFI) of each TF. (C, E) Plots are gated on WT or *Batf*^-/-^ P14 cells. (F) Expression of BATF and Tim-3 in WT P14 cells (dot plot) and Tim-3 expression in BATF^hi^ and BATF^lo^ WT P14 cells (bar plot) derived from spleen of the recipient mice in A. \*\*\*\**p* < 0.0001 (unpaired Student’s *t*-test). Data are representative of two independent experiments with four to five mice in each experiment.

### BATF regulates gene expression and chromatin remodeling of CD8^+^ T cells during early effector differentiation

We next sought to investigate the mechanisms by which BATF mediates the differentiation of cytolytic effector CD8^+^ T cells in detail. Previous studies have shown that EEC differentiate into cytolytic SLEC or MPEC at around 4-5 days post infection (Delpoux et al., 2021; Joshi et al., 2007; Obar & Lefrançois, 2010), and that *Batf*^-/-^ CD8^+^ T cells exhibited a proliferative defect at 4 days post-infection (Kurachi et al., 2014). To assess whether BATF programs cell fate determination preceding SLEC/MPEC differentiation occurs, therefore, we analyzed WT and *Batf*^-/-^ P14 cells at 3 days post-infection. Naive WT recipients were adoptively transferred with WT and *Batf*^-/-^ P14 cells and infected with LCMV Arm as indicated in Supplementary figure 1A. At 3 days post-infection, effector P14 cells were sorted and gene expression profiles were analyzed by RNA-seq. We detected a slight reduction in the frequency of *Batf*^-/-^ P14 cells at this time point (Supplementary figure 2A). We observed that the lack of BATF resulted in profound alterations in gene expression, with 3,835 genes downregulated and 3,820 genes upregulated (Fig. 2A). Principal component analysis (PCA) further delineated that WT and *Batf*^-/-^ effector P14 cells underwent distinct transcriptomic programs in response to LCMV infection, whereas naive WT and *Batf*^-/-^ P14 cells clustered together (Fig. 2B). While the extent to which the numbers of genes, whose expression changed during the transition from naive state, was comparable between WT and *Batf*^-/-^ P14 cells, we found that *Batf*^-/-^ P14 cells underwent dysregulated transcriptional programs in response to LCMV infection, with only 4.2% of upregulated genes and 4.4% of downregulated genes observed in WT P14 cells being shared by *Batf*^-/-^ P14 cells (Supplementary figure 2B). As expected, gene set enrichment analysis (GSEA) revealed a significant enrichment of genes associated with SLEC-like and other effector CD8^+^ T cell phenotypes in WT P14 cells, whereas *Batf*^-/-^ P14 cells exhibited transcriptomic signatures representing MPEC and naive cells (Fig. 2C, Supplementary fig. 2C). Correspondingly, *Batf*-deficient cells displayed reduced expression of genes related to SLEC differentiation (*Tbx21*, *Ezh2*), while the expression of genes characteristic of naive or memory CD8^+^ T cells, such as *Tcf7*, *Foxo1*, *Sell*, *Cd127* were higher in *Batf*^-/-^ populations (Supplementary figure 2D, E). In addition, genes associated with nucleic acid metabolism and cell cycle regulation were predominantly found in WT P14 cells (supplementary figure 2C). We also noted that genes involved in apoptotic pathways were predominantly enriched in *Batf*^-/-^ effector P14 cells (Supplementary table 1), corresponding to the previous study that showed increased expression of cell death markers in *Batf*^-/-^ effector CD8^+^ T cells (Kurachi et al., 2014). Consistently, the expression of a pro-apoptotic gene *Bcl2l11* (encoding Bim), which is regulated by BATF in CD4^+^ T cells (Titcombe et al., 2023), was significantly higher in *Batf*^-/-^ cells (Supplementary figure 2F).

**Fig. 2.**
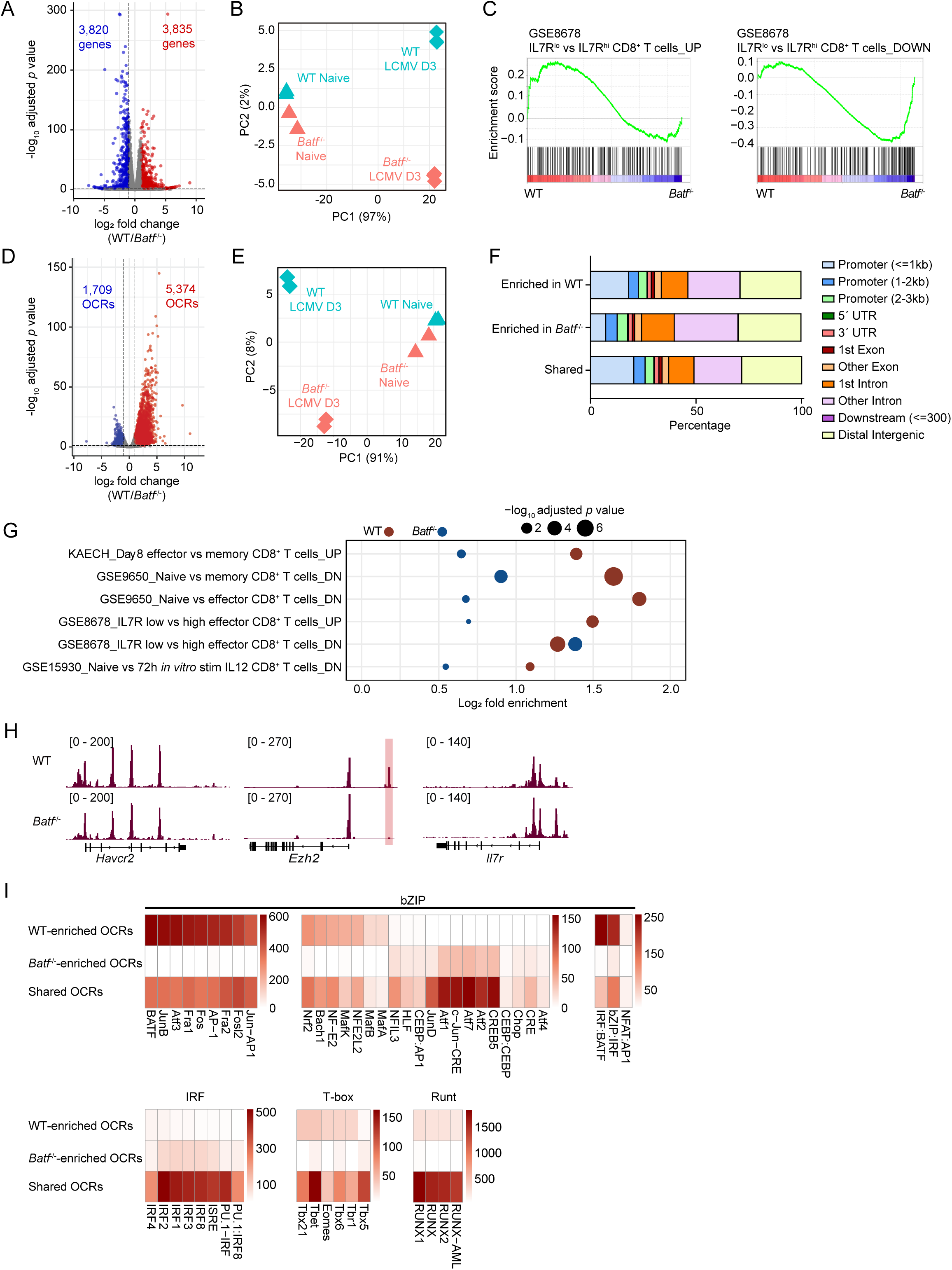
BATF regulates transcriptomic and epigenomic reprogramming of CD8^+^ T cells during early effector differentiation. WT recipient mice were adoptively transferred with congenically distinct WT and *Batf*^-/-^ P14 cells mixed at 1:1 ratio (1 × 10^6^ each cell), followed by infection of the recipient mice with LCMV Arm. WT and *Batf*^-/-^ P14 cells collected from spleen of the recipient mice at day 3 post-infection (“effector”) and naive WT and *Batf*^-/-^ P14 cells were subjected to RNA-seq (effector, *n* = 3; naive, *n* = 2) and ATAC-seq (*n* = 2). (A) Volcano plot showing differences in gene expression between WT and *Batf*^-/-^ effector P14 cells. (B) Principal component analysis (PCA) of RNA-seq data of naive and effector P14 cells. (C) Gene set enrichment analysis (GSEA) between WT and *Batf*^-/-^ effector cells of IL-7R^lo^ short-lived effector cells (SLEC) signature (left) and IL-7R^hi^ memory-precursor effector cells (MPEC) signature (right) (GSE8678). (D) Volcano plot represents differences in chromatin accessibility between WT and *Batf*^-/-^ effector P14 cells. (E) PCA of ATAC-seq data of naive and effector P14 cells. (F) Peak annotation of open chromatin regions (OCRs) differentially accessible or shared between WT and *Batf*^-/-^ effector P14 cells. (G) Fold enrichment for selected CD8^+^ T cells-related immunologic signatures of OCRs differentially accessible between WT and *Batf*^-/-^ effector P14 cells analyzed using GSEA C7 immunological signature gene sets and rGREAT. Gene sets associated with WT-enriched OCRs are shown in red and gene sets associated with *Batf*^-/-^ -enriched OCRs are shown in blue. (H) Representative ATAC-seq signal tracks for WT and *Batf*^-/-^ effector P14 cells. (I) Enrichment of selected TF motifs in differentially accessible or shared OCRs between WT and *Batf*^-/-^ effector P14 cells. Color bars represent enrichment scores (-log_10_ *p*-value).

BATF is critical for the chromatin remodeling in response to antigenic stimulation in T cells (Pham et al., 2019; Tsao et al., 2022). Therefore, to examine the role of BATF in chromatin remodeling during early effector differentiation of CD8^+^ T cells, we next analyzed chromatin accessibility of naive and effector WT and *Batf*^-/-^ P14 cells by assay for transposase-accessible chromatin sequencing (ATAC-seq). We identified 111,415 open chromatin regions (OCRs) across all samples analyzed, among which 5,374 regions exhibited increased accessibility in WT effector P14 cells, whereas 1,709 regions were less accessible (Fig. 2D). PCA revealed distinct chromatin landscapes for WT and *Batf*^-/-^ P14 cells in LCMV-infected mice, while naive WT and *Batf*^-/-^ P14 cells co-localized (Fig. 2E). When OCRs gained or lost their accessibility during transition from naive cells were examined, 53% of gained OCRs and 72% of lost OCRs in WT P14 cells were shared by *Batf*^-/-^ P14 cells (Supplementary figure 3A). We next analyzed OCRs differentially accessible (fold-change < -2.0 or > 2.0, and FDR < 0.05) and those shared by effector WT and *Batf*^-/-^ P14 cells (Supplementary figure 3B). Peak annotations revealed that OCRs enriched in *Batf*^-/-^ P14 cells were less frequently observed at promoter regions within 1 kb from the transcription start site (TSS), whereas WT-enriched and shared OCRs displayed similar frequencies in each annotation (Fig. 2F). To further investigate the relationship between chromatin landscapes and transcriptional programs, we performed functional enrichment analysis for OCRs enriched in WT and *Batf*^-/-^ P14 cells, with rGREAT (Gu & Hübschmann, 2023; McLean et al., 2010). OCRs proximal to the genes highly expressed in effector, activated, and SLEC populations compared with naive, memory, and MPEC were more frequently observed in WT P14 cells, while OCRs proximal to the genes representing MPEC-like signatures were enriched in *Batf*^-/-^ P14 cells (Fig. 2G). Correspondingly, WT P14 cells exhibited increased accessibilities at several genes related to effector properties, including *Havcr2*, *Cx3cr1*, and *Maf* compared to *Batf*^-/-^ P14 cells, while accessibilities at some genes highly expressed in naive or memory populations (*Il7r*, *Ccr7*, *Klf2*) decreased (Fig. 2H, Supplementary figure 3C). In addition, the accessibility at the enhancer region of *Ezh2* gene (Ochiai et al., 2018) was almost completely abolished in the absence of BATF (Fig. 2H, highlighted).

We next performed TF motif enrichment analysis on the differentially accessible and shared OCRs. We observed that several AP-1 (bZIP) motifs, including BATF and JunB, were predominantly found in WT-enriched OCRs (Fig. 2I). Notably, certain AP-1 motifs, such as JunD and c-Jun, displayed higher enrichment scores in *Batf*^-/-^-enriched OCRs. Importantly, WT-enriched OCRs exhibited significant enrichment of IRF:BATF and bZIP:BATF motifs compared to both *Batf*^-/-^-enriched and shared OCRs, while the enrichment of NFAT:AP-1 motif was similar between WT-enriched and shared OCRs. However, IRF motifs were more prevalent in OCRs enriched in *Batf*^-/-^ rather than WT-enriched OCRs. The cooperation between BATF and IRFs, especially IRF4, at AP-1-IRF composite DNA elements (AICEs) is critical for differentiation of Th17 cells (Ciofani et al., 2012; Li et al., 2012; Murphy et al., 2013). These findings indicate that BATF remodels chromatin landscapes and recruits IRFs to AICEs to mount an effector response of CD8^+^ T cells. We also found that motifs of T-box and Runt, which regulate effector differentiation of CD8^+^ T cells (Joshi et al., 2007; Shan et al., 2017), were more enriched in WT-enriched OCRs, whereas ETS and Zf motifs were predominantly enriched in shared OCRs (Supplementary figure 3D). Taken together, these data suggest that BATF plays a central role in establishing epigenomic and transcriptomic landscapes to initiate the differentiation of cytolytic effector CD8^+^ T cells.

### BATF orchestrate the binding of transcription factors to DNA during effector CD8^+^ T cell differentiation

Given that chromatin remodeling during effector CD8^+^ T cells differentiation was highly dependent on BATF, we hypothesize that the accessibilities of TFs to DNA are also BATF-dependent. To assess this, we examined the bindings of TFs to DNA in effector WT and *Batf*^-/-^ P14 cells at days 3 after LCMV Arm infection by cleavage under target & release using nuclease (CUT&RUN) assay (Skene & Henikoff, 2017). We analyzed T-bet as a representative TF which mediates SLEC differentiation (Joshi et al., 2007), JunB and JunD as AP-1 transcription factors that may form heterodimers with BATF (Li et al., 2012; Tussiwand et al., 2012), and IRF4, a crucial partner TF of BATF (Ciofani et al., 2012; Iwata et al., 2017; Li et al., 2012; Murphy et al., 2013; Seo et al., 2021; Tussiwand et al., 2012). In addition, we examined the binding of FoxO1, a TF programs memory commitment (Delpoux et al., 2017; Michelini et al., 2013), as it reciprocally inhibits BATF in CD8^+^ T cells (Delpoux et al., 2021). We found drastic changes in the binding profiles of each TF in the absence of BATF (Fig. 3A, Supplementary figure 4A), with a significant reduction in the numbers of peaks (Supplementary figure 4B). We observed a substantial reduction of TF bindings at the enhancer region of *Ezh2* in *Batf*^-/-^ cells, consistent with the diminished chromatin accessibility at this genomic locus (Supplementary figure 4A and Fig. 2H, highlighted). Differential binding analysis revealed profound loss of bindings for IRF4 and T-bet in *Batf*^-/-^ populations, implicating their dependency on BATF for binding (Fig. 3B). In particular, the binding of IRF4 was predominantly enriched in WT P14 cells, with over 75% of IRF4-bound loci co-bound by BATF in WT P14 cells (Figs. 3B, C). JunD also displayed significant overlap with BATF in WT P14 cells; however, its binding was significantly enriched in *Batf*^-/-^ P14 cells. Moreover, over 40% of JunD-bound loci in *Batf*^-/-^ P14 cells were uniquely found in *Batf*^-/-^ samples, whereas those of other TFs were largely observed in WT samples as well (Fig. 3D, Supplementary figure 4C). In addition, over 85% of JunD-bound loci co-localized with JunB in *Batf*^-/-^ cells, indicating the increase in JunB-JunD heterodimerization in the absence of BATF (Supplementary figure 4D). These results suggest that BATF interacts with JunD to control its proper binding. We also found that approximately 94% of BATF-bound loci co-localized by JunB-bound regions (Fig. 3E), suggesting that the cooperation with JunB is crucial for BATF-mediated effector CD8^+^ T cell differentiation, as previously reported (Chang et al., 2018; Murphy et al., 2013). Notably, bindings of T-bet were also frequently detected in close proximity to BATF-bound loci; however, T-bet-bound loci co-bound by BATF remained 25%, and we observed binding patterns of T-bet were less correlated with BATF, compared to those of JunB and JunD (Fig. 3F).

**Fig. 3.**
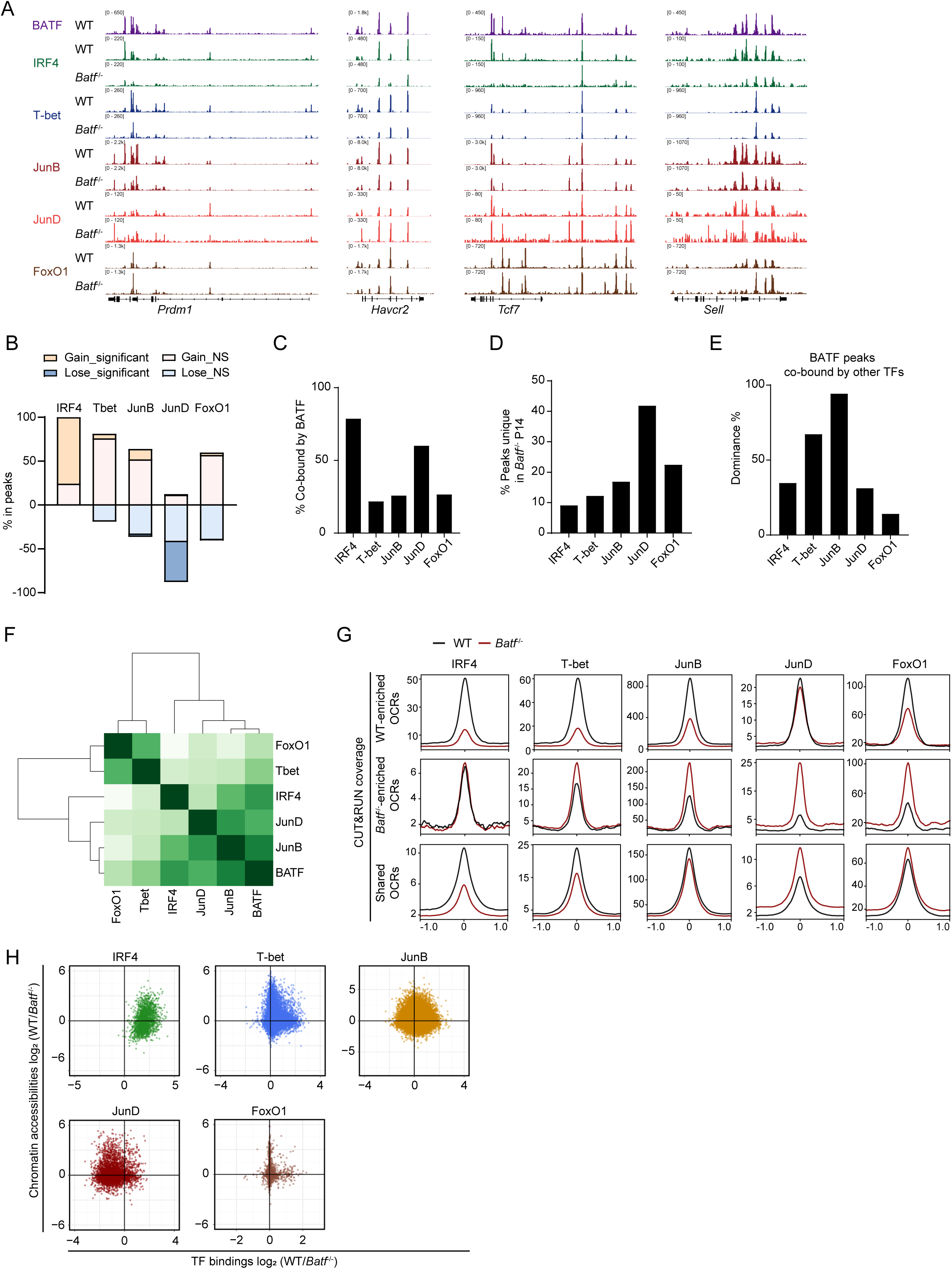
BATF orchestrates the bindings of key transcription factors during effector CD8^+^ T cell differentiation. WT recipient mice were adoptively transferred with congenically distinct WT and *Batf*^-/-^ P14 cells mixed at 1:1 ratio (1 × 10^6^ each cell), followed by infection of the recipient mice with LCMV Arm. WT and *Batf*^-/-^ effector P14 cells were collected from spleen of the recipient mice at day 3 post-infection and subjected to CUT&RUN (*n* = 2) analysis. (A) Representative CUT&RUN signal tracks for WT and *Batf*^-/-^ effector P14 cells. (B) Differences in TF bindings between WT and *Batf*^-/-^ effector P14 cells. Peaks were considered as differentially bound region if they have a false-discovery rate (FDR) < 0.05. (C, D) Percentage of peaks in each TF co-bound by BATF in WT P14 cells (C) and uniquely found in *Batf*^-/-^ P14 cells (D). (E) Percentages of BATF peaks co-bound by each TF. (F) Heatmap represents correlations between the peaks of each TF in WT P14 cells. (G) CUT&RUN signal coverages at OCRs differentially accessible or shared between WT and *Batf*^-/-^ effector P14 cells in Fig. 2. Horizontal axis represents distance from peak center. (H) Scatter plots showing relationship between differences in TF bindings (horizontal axis) and differences in chromatin accessibilities (vertical axis) between WT and *Batf*^-/-^ effector P14 cells. (A, C-E, G) Data from biological replicates were merged and analyzed.

Next, we integrated CUT&RUN and ATAC-seq data to further assess the effect of BATF deficiency on TF bindings (Figs. 3G, H and Supplementary figure 4E). We found that IRF4 bindings were significantly enriched in WT P14 cells, both in WT-enriched OCRs and shared OCRs, suggesting that IRF4 bindings are dependent on the cooperation with BATF rather than chromatin accessibility. Conversely, genomic regions bound by T-bet, JunB, and FoxO1 largely corresponded to OCRs, indicating the dependency of their bindings on chromatin accessibility. Remarkably, bindings of JunD in WT-enriched OCRs was equivalent between WT and *Batf*^-/-^ cells, whereas those in *Batf*^-/-^-enriched OCRs and shared OCRs were predominantly observed in *Batf*^-/-^ cells, implicating that BATF negatively regulates JunD binding. Correctively, our data demonstrated that BATF organizes the bindings of other transcription factors and coordinates epigenomic programs during early effector CD8^+^ T cell differentiation.

### IRF4 controls BATF-mediated epigenetic programs during early effector differentiation of CD8^+^ T cells

We showed that BATF is crucial for the binding of IRF4 during effector differentiation of CD8^+^ T cells. Next, we examined whether BATF and IRF4 are reciprocally dependent. We crossed *Irf4*-floxed mice with CD4^Cre^ P14 mice to generate P14 mice that deficient in IRF4 expression in the CD4^+^ T cells and CD8^+^ T cells (hereafter IRF4cKO mice). Congenically distinct IRF4cKO and CD4^Cre^ P14 cells were adoptively transferred at a ratio of 1:1 into naive WT recipients, and recipient mice were infected with LCMV Arm. At day 3 post-infection, IRF4cKO effector P14 cells exhibited impaired expansion (Supplementary figures 5A and B), as observed in *Batf*^-/-^ effector P14 cells (Supplementary figure 2A). We observed that the expression of TCF-1 was elevated in IRF4cKO cells, while those of BATF and T-bet were higher in CD4^Cre^ populations (Supplementary figure 5C). Next, we analyzed chromatin accessibilities of CD4^Cre^ and IRF4cKO effector P14 cells and found that IRF4-deficient cells were in distinct chromatin landscapes from CD4^Cre^ cells (Figure 4A, Supplementary figure 6A). When differentially accessible regions between each sample was analyzed, OCRs enriched in IRF4cKO P14 cells were less frequently observed at promoter regions proximal to TSS, a trend like that observed in BATF-deficient cells (Supplementary figure 6B, Fig. 2F). In addition, we observed a significant consistency in the effects of IRF4 loss on chromatin accessibilities with those of BATF loss (Fig. 4B and Supplementary figure 6C), as reported previously (Tsao et al., 2022). By PCA, in contrast, *Batf*^-/-^ and IRF4cKO effector P14 cells localized at distinct principal component space (Fig. 4C). Furthermore, IRF4cKO effector P14 displayed a considerable increase in chromatin accessibility compared to *Batf*^-/-^ effector cells (Supplementary figure 8D), suggesting that although BATF and IRF4 function cooperatively, they also play distinct roles in remodeling chromatin landscapes.

**Fig. 4.**
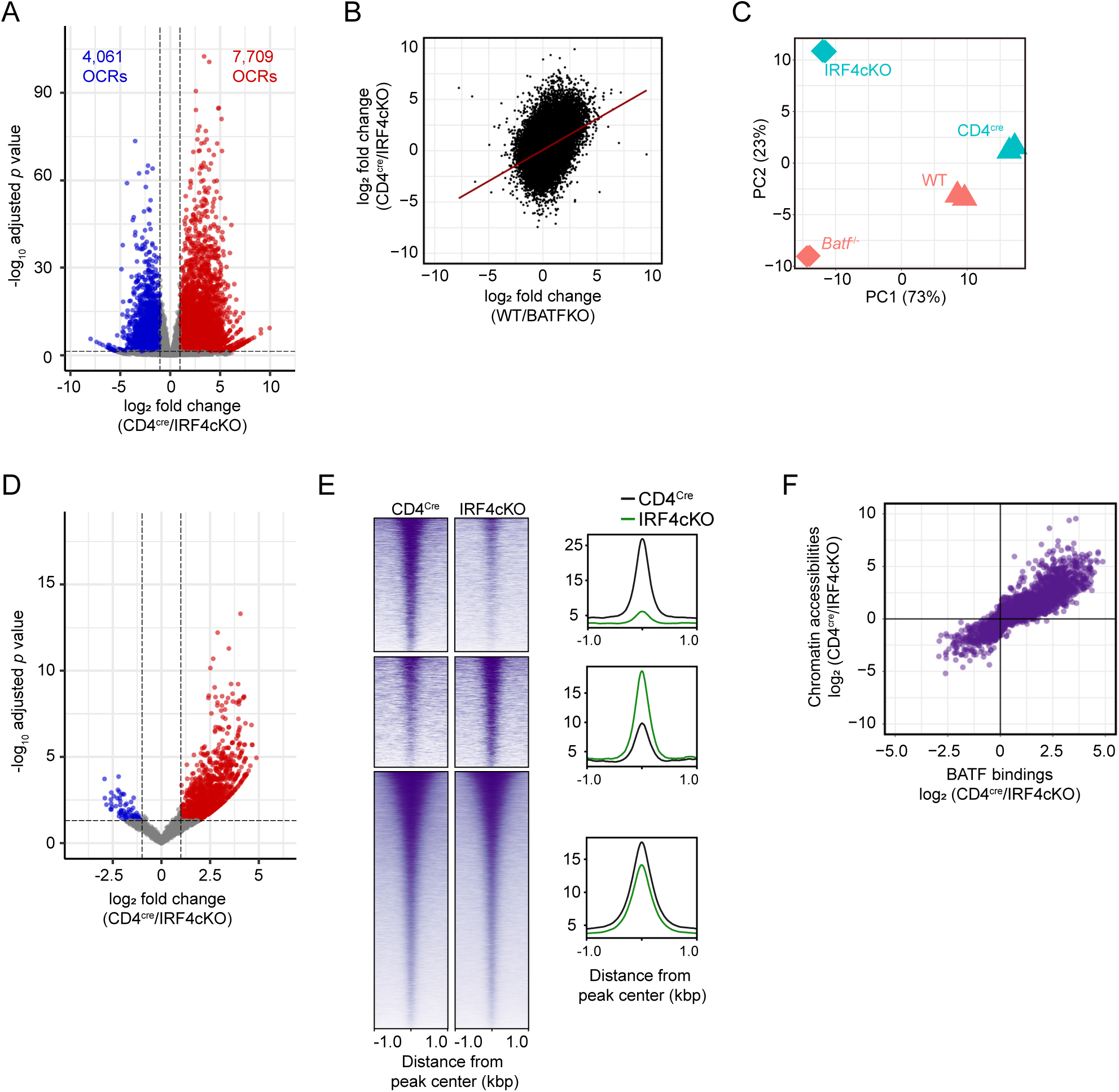
IRF4 is indispensable for the chromatin remodeling but not for BATF binding during effector differentiation of CD8^+^ T cells. WT recipient mice were adoptively transferred with congenically distinct CD4^Cre^ and IRF4cKO P14 cells mixed at 1:1 ratio (1 × 10^6^ each cell), followed by infection of the recipient mice with LCMV Arm. CD4^Cre^ and IRF4cKO effector P14 cells were collected from spleen of the recipient mice at day 3 post-infection and subjected to ATAC-seq and BATF CUT&RUN analyses (*n* = 2). (A) Volcano plot represents differences in chromatin accessibility between CD4^Cre^ and IRF4cKO effector P14 cells. (B) Scatter plots showing relationship between differences in chromatin accessibilities between WT and *Batf*^-/-^ effector P14 cells (horizontal axis) in Fig. 2 and between CD4^Cre^ and IRF4cKO effector P14 cells (vertical axis). (C) PCA of ATAC-seq data of WT, *Batf*^-/-^ effector P14 cells in Fig. 2 and CD4^Cre^ and IRF4cKO effector P14 cells. (D) Volcano plot represents differences in the bindings of BATF to DNA between CD4^Cre^ and IRF4cKO effector P14 cells. (E) BATF CUT&RUN signal coverages at OCRs differentially accessible or shared between CD4^Cre^ and IRF4cKO effector P14 cells. Horizontal axis represents distance from peak center. (F) Scatter plots showing relationship between differences in BATF bindings (horizontal axis) and differences in chromatin accessibilities (vertical axis) between CD4^Cre^ and IRF4cKO effector P14 cells.

To assess whether the loss of IRF4 impacts the binding of BATF to DNA, we next performed CUT&RUN using CD4^Cre^ and IRF4cKO cells to analyze the accessibility of BATF. We found a substantial loss of BATF binding in the absence of IRF4 (Figure 4D, Supplementary figures 6E, F). However, when CUT&RUN and ATAC-seq data were integrated, we observed a significant correlation between BATF binding and chromatin accessibility (Figs. 4E, F). These results indicated that the binding of BATF was predominantly dependent on chromatin accessibility. Collectively, our data proposed that IRF4 remodels chromatin landscapes of CD8^+^ T cells during effector differentiation like BATF does, whilst IRF4 may not be required for the binding of BATF to DNA.

### BATF-IRF4 interaction is indispensable for effector CD8^+^ T cell differentiation

Previous studies have reported that BATF interact with IRF4 at its leucin zipper motif (histidine-55 residue), and that BATF-IRF4 interaction is critical for Th17 differentiation of CD4^+^ T cells and dendritic cell development(Ciofani et al., 2012; Li et al., 2012; Murphy et al., 2013; Tussiwand et al., 2012). In addition, BATF counteracts exhaustion of tumor-infiltrating CAR T cells through the cooperation with IRF4 (Seo et al., 2021). However, the requirement of BATF-IRF4 interaction during effector CD8^+^ T cell differentiation have not been studied. To address this, WT and *Batf*^-/-^ P14 cells were retrovirally transduced with BATF or BATF H55Q mutant (BATF H55 was replaced by glutamine residue) and adoptively transferred into infection-matched recipient mice (Supplementary figures 7A, B). Consistent with the previous study (Kurachi et al., 2014), reconstitution of BATF into *Batf*^-/-^ P14 cells reinvigorated their proliferative potential, whereas overexpression of BATF into WT P14 cells influenced the expansion of P14 cells (Fig. 5A, Supplementary figures 7C, E, F). However, transduction of BATF H55Q mutant could not restore the clonal expansion of *Batf*^-/-^ P14 cells, indicating that interaction with IRF4 is critical for BATF-mediated effector response of CD8^+^ T cells. Remarkably, WT P14 cells transduced with BATF H55Q mutant failed in proliferation, suggesting that BATF H55Q mutant has a dominant-negative effect on the endogenous BATF. We also found that P14 cells transduced with BATF predominantly differentiated into KLRG1^hi^CD127^lo^ SLEC population, while considerable increase of KLRG1^lo^CD127^hi^ MPECs was observed in cells transduced with empty RV or H55Q mutant (Fig. 5B and Supplementary figures 7D, G, H). It has been reported that phosphorylation of serine-43 residue of BATF is critical for its binding to DNA, with the constitutively non-phosphorylated S43A (S43 was replaced by alanine residue) mutant potentially increasing DNA binding, whereas constitutively phosphorylated S43D (S43 was replaced by aspartic acid residue) mutant shows decreased DNA binding (Deppmann et al., 2003). We observed that the proliferation and phenotype of both WT and *Batf*^-/-^ P14 cells transduced with S43A mutant was comparable with those transduced with BATF, suggesting that binding intensity of BATF sufficient to initiate effector differentiation of CD8^+^ T cells. Meantime, *Batf*^-/-^ P14 cells transduced with BATF S43D mutant could not proliferate in response to LCMV infection and exhibited higher frequency of MPEC populations, demonstrating that the binding to DNA is critical for BATF-mediated effector differentiation. Of note, WT P14 cells expressing S43D mutant exhibited an impaired proliferation and increase in MPEC, indicating that S43D mutant also has a dominant-negative effect on endogenous BATF, while its magnitude seemed lower than that of H55Q mutant. Notably, transduction of BATF H55Q-S43A double mutant did not initiate clonal expansion of *Batf*^-/-^ P14 cells and showed a dominant-negative effect on WT P14 cells, like BATF H55Q single mutant. These data further suggest that interaction with IRF4 via BATF H55 residue, rather than the intensity of BATF binding, is critical BATF-mediated effector differentiation of CD8^+^ T cells.

**Fig. 5.**
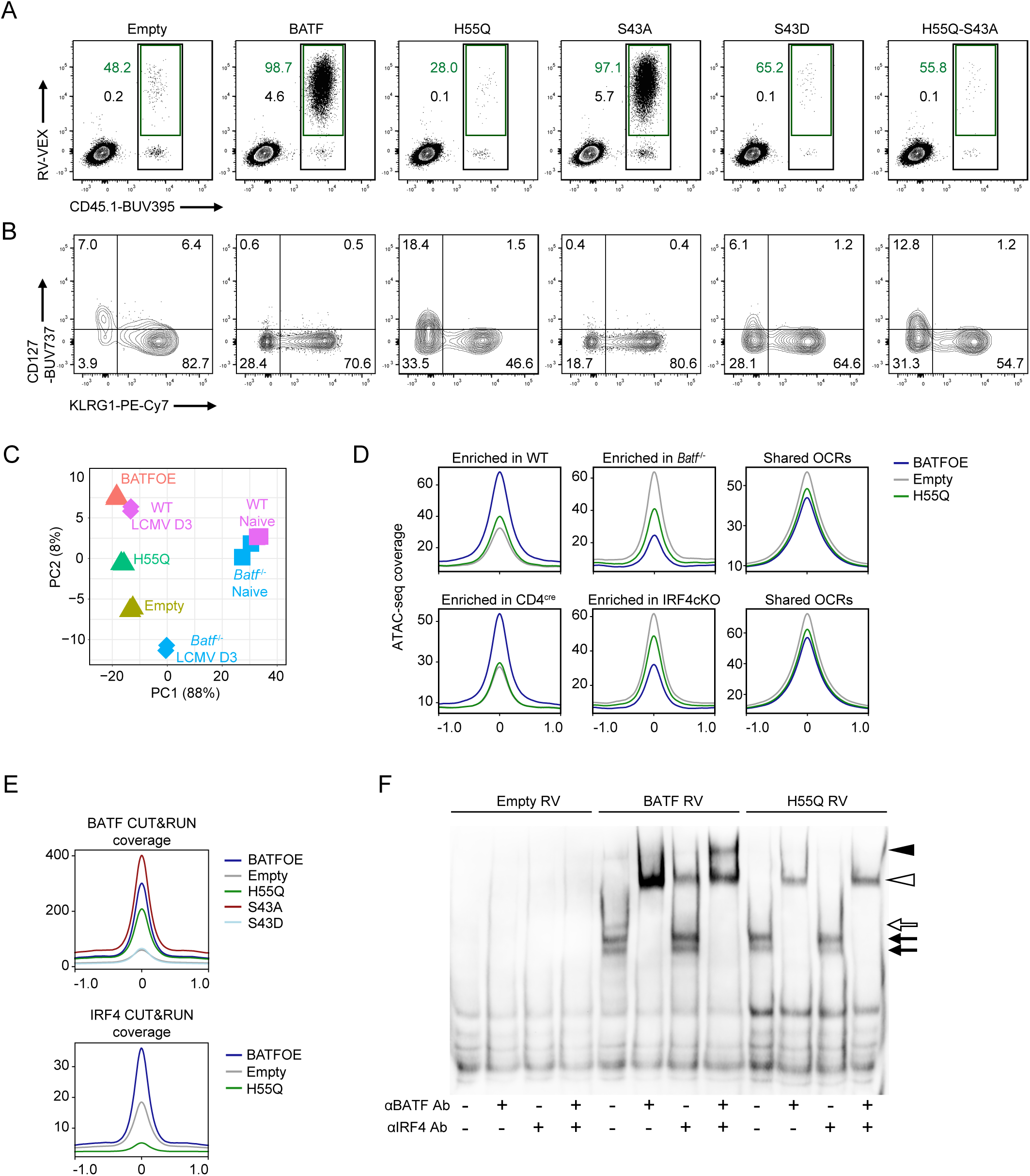
Direct interaction between BATF and IRF4 is critical for effector differentiation of CD8^+^ T cells. (A, B) Flow cytometric analysis of splenocytes of WT recipient mice infected with LCMV Arm, and adoptively transferred with WT or *Batf*^-/-^ P14 cells (1 × 10^5^ each cell) transduced with empty retrovirus (RV) or RV overexpressing *Batf*, *Batf* H55Q, *Batf* S43A, *Batf* S43D, or *Batf* H55Q-S43A after 1 d stimulation *in vitro* with anti-CD3 and anti-CD28 antibodies (Supplementary figure 7), assessed at day 8 post-infection of recipient mice. Data are representative of two independent experiments with three to five mice in each experiment. (A) Plots are gated on total CD8^+^ T cells and numbers indicate percent *Batf*^-/-^ P14 cells among total CD8^+^ T cells (black) and percent RV-transduced VEX^+^ cells among total *Batf*^-/-^ P14 cells (green). (B) Expression of KLRG1 and CD127 on RV-transduced *Batf*^-/-^ P14 cells; plots are gated on VEX^+^ P14 cells. (C, D) *Batf*^-/-^ effector P14 cells transduced with empty RV, RV overexpressing *Batf* (BATFOE) or *Batf* H55Q mutant were collected from spleen of the recipient mice at day 3 post-infection and subjected to ATAC-seq analysis (*n* = 2). (C) PCA of ATAC-seq data of *Batf*^-/-^ effector P14 cells transduced with indicated RV, and that of WT and *Batf*^-/-^ effector P14 cells in Fig. 2. (D) ATAC-seq signal coverages of *Batf*^-/-^ effector P14 cells transduced with indicated RV at OCRs differentially accessible or shared between WT and *Batf*^-/-^ P14 cells in Fig. 2 (top), or between CD4^Cre^ and IRF4cKO effector P14 cells in Fig. 4 (bottom). (E, F) *Batf*^-/-^ P14 cells transduced with empty RV, RV overexpressing *Batf*, *Batf* H55Q, S43A, or S43D mutant were cultured *in vitro*, and subjected to CUT&RUN (E) or EMSA (G) at day 3 after RV transduction. (E) BATF (upper) or IRF4 (lower) CUT&RUN signal coverage of *Batf*^-/-^ P14 cells transduced with indicated RV (*n* = 2). Data from biological replicates were merged and analyzed. (F) Nuclear extracts of P14 cells transduced with indicated RV were analyzed using the AICE1 probe (Tussiwand et al., 2012), in the presence or absence of antibodies against BATF and IRF4 as indicated. Data is representative of two independent experiments.

We next examined chromatin accessibility of *Batf*^-/-^ P14 cells transduced with BATF or BATF H55Q mutant at days 4 post-infection (at days 3 after adoptive transfer). As expected, *Batf* RV-transduced P14 cells showed a significant increase in chromatin accessibility compared to empty RV-transduced cells, supporting the critical role of BATF in the chromatin remodeling during early effector response of CD8^+^ T cells (Supplementary figure 8A). In contrast, P14 cells transduced with H55Q RV failed in proper epigenomic modification, as represented by substantial decrease in chromatin accessibility compared to *Batf* RV-transduced cells. Thus, BATF-mediated epigenomic reprogramming during effector differentiation requires a cooperation with IRF4. Meanwhile, PCA revealed that H55Q RV-transduced cells localized at a principal component space closer to BATF-expressing cells compared to empty RV-transduced cells, suggesting an IRF4-independent role for BATF in chromatin remodeling (Fig. 5C). Consistently, chromatin accessibility of H55Q-expressing cells slightly increased compared to that of empty RV-transduced cells (Supplementary figure 8A). Moreover, while empty RV- and H55Q RV-transduced cells displayed chromatin accessibilities largely corresponding to those of *Batf*^-/-^ P14 cells (Supplementary figure 8B, highlighted in red), we also identified loci whose accessibilities in H55Q-expressing cells were equivalent to those of WT and *Batf* RV-transduced *Batf*^-/-^ P14 cells (highlighted in blue). Motif enrichment analysis revealed that motifs of several AP-1 TFs including BATF and JunB, and IRF:AP-1 motifs were more enriched in OCRs specific to BATF-expressing cells compared to OCRs specific to H55Q-expressing cells. Conversely, JunD and cJun motifs were predominantly enriched in H55Q-expressing cells (Supplementary figure 8C). Strikingly, ATAC-seq coverage of H55Q-expressing cells at OCRs enriched in *Batf*^-/-^ cells were intermediate between cells transduced with empty RV and *Batf* RV, while chromatin accessibility at IRF4cKO-enriched OCRs were more comparable between empty RV-transduced cells and H55Q expressing cells (Fig. 5D). These results further indicate that BATF regulates chromatin accessibility of CD8^+^ T cells in IRF4-dependent and independent manners.

Next, we sought to corroborate that BATF binds to DNA in an IRF4-independent manner, as indicated in Fig. 4. *Batf*^-/-^ P14 cells were transduced with empty RV or RV overexpressing *Batf*, *Batf* H55Q, *Batf* S43A, *Batf* S43D and cultured *in vitro*. RV^+^ cells were sorted 3 days after transduction, and the bindings of each BATF protein were analyzed by CUT&RUN. As expected, S43D mutant lost its binding to DNA, whereas S43A mutant showed an increased signal compared to wild-type BATF (Fig. 5E, Supplementary figure 8D). Importantly, while H55Q mutant displayed reduced bindings in comparison to BATF, it retained binding capacity to DNA. Peak annotations revealed that the binding of H55Q mutant was more enriched for promoters within 1 kb from TSS, whereas wild-type BATF and S43A mutant showed similar binding patterns (Supplementary figure 8E). Of note, we identified several loci at which H55Q mutant bound more strongly than BATF and S43A mutant (Supplementary figure 8D, highlighted in red). Motif enrichment of peaks specific to wild-type BATF-expressing cells or H55Q-expressing cells as well as shared peaks revealed an enrichment of motifs bound by AP-1 transcription factors. Notably, while the motif bound by BATF-IRF complex exhibited a significant enrichment in peaks specific to wild-type BATF and to a lesser extent in shared peaks, we did not detect an enrichment of the IRF:BATF motif in H55Q-specific peaks (Supplementary table 2). When BATF CUT&RUN signals were integrated to ATAC-seq data, we found the correlation between BATF binding and chromatin accessibility (Supplementary figure. 8F). These data support our hypothesis that binding of BATF to DNA primarily depends on chromatin accessibility rather than the interaction with IRF4. We also profiled the binding of IRF4 in cells transduced with empty RV, *Batf* RV, or H55Q RV. As expected, IRF4 binding was drastically reduced in empty- and H55Q RV-transduced cells, indicating that the interaction with BATF is indispensable for the binding of IRF4 (Fig. 5E, Supplementary figures 8D and F). Intriguingly, IRF4 CUT&RUN coverage in H55Q-expressing cells were almost completely abolished, and substantially lower than that in empty RV-transduced cells. Motif enrichment analysis of peaks specific to or shared between each cell revealed a significant enrichment of AP-1 and IRF:BATF motifs in peaks specific to BATF-expressing cells and shared peaks; however, peaks specific to H55Q-expressing cells exhibited an enrichment of zinc finger transcription factors including Krüppel-like factors, and that of AP-1 factors was not detected (Supplementary table 3). Integrative analysis with ATAC-seq data demonstrated that the binding of IRF4 in empty RV-transduced cells markedly decreased at OCRs enriched in *Batf* RV-transduced cell and shared OCRs compared to those in BATF-expressing cells, whereas the CUT&RUN coverage at H55Q-enrched OCRs was comparable between these two samples (Supplementary figure 8E). These results ascertained that IRF4 binding is exhaustively dependent on BATF. We finally performed electrophoretic mobility shift assays (EMSA) to biochemically corroborate the interaction between BATF and IRF4. Nuclear extracted from *Batf*^-/-^ P14 cells transduced with empty RV, *Batf* RV, or H55Q RV were incubated with the AICE consensus probe, in the presence or absence of antibodies against BATF and IRF4. No signal was detected in empty RV-transduced samples, indicating that IRF4 binding to the AICE probe necessitates BATF. In P14 cells transduced with *Batf* RV and H55Q RV, signals representing BATF complexes were observed (white and black arrows), which underwent supershift upon incubation with an anti-BATF antibody (white arrowhead). These findings demonstrated that both wild-type BATF and the H55Q mutant are capable of binding to the AICE probe, as previously demonstrated in dendritic cells and CD4^+^ T cells (Glasmacher et al., 2012; Li et al., 2012; Murphy et al., 2013; Tussiwand et al., 2012). Notably, a distinct signal was identified in *Batf* RV-transduced cells (white arrow), which underwent supershift upon treatment with antibodies against BATF and IRF4 (black and white arrowheads). Thus, BATF and IRF4 cooperatively bind to AICE in CD8^+^ T cells. Conversely, in H55Q RV-transduced cells, no supershift was observed with an anti-IRF4 antibody. Collectively, these results manifested that the direct interaction between BATF, mediated by the H55 residue of BATF, is essential for IRF4 binding to DNA, as indicated by our genomic analyses. Taken together, these results suggested that BATF initiates the chromatin reorganization and recruits IRF4 to AICEs, subsequently triggering dynamic modifications of epigenomic and transcriptomic landscapes, thereby enabling CD8^+^ T cells to undergo effector differentiation programs.

## Discussion

Mounting studies have revealed that BATF regulates the differentiation and function of lymphocytes in cooperation with IRF4, through the modification of epigenomic and transcriptomic landscapes (Y. Chen et al., 2021; Ciofani et al., 2012; Ise et al., 2011; Itahashi et al., 2022; Kurachi et al., 2014; Murphy et al., 2013; Pham et al., 2019; Tsao et al., 2022). In addition, previous studies demonstrated that loss of BATF perturbs the bindings of several transcription factors *in vitro* (Pham et al., 2023; Tsao et al., 2022); however, detailed mechanisms how BATF facilitates effector CD8^+^ T cell differentiation *in vivo* have remained poorly understood. In the present study, we investigated epigenomic and transcriptomic events mediated by BATF during early effector differentiation of CD8^+^ T cells *in vivo* and found that BATF initiates the configuration of epigenomic landscapes and orchestrate the deployments of key TFs directly (IRF4 and AP-1) or indirectly (T-bet, FoxO1) in response to acute viral infection.

Most importantly, we demonstrated that the direct interaction with IRF4 is essential for BATF-mediated effector differentiation of CD8^+^ T cells, and that IRF4 exclusively depends on BATF for its binding to genome. Several studies have indicated that the binding of IRF4 to AICEs requires BATF (Glasmacher et al., 2012; Murphy et al., 2013; Tussiwand et al., 2012); meanwhile, we observed that IRF4 showed a genome-wide loss of binding in the absence of BATF, despite the IRF4 motif was similarly observed at OCRs in both WT and *Batf*^-/-^ effector P14 cells. Previous studies have demonstrated the low affinity of IRF4 to DNA and hence, heterodimerization with partner TFs, such as BATF, NFATc2, STAT3, STAT6, and PU.1, is critical for IRF4 to bind to DNA (Brass et al., 1999; Gupta et al., 1999; Kwon et al., 2009; Li et al., 2012; Rengarajan et al., 2002). In addition, IRF4 can bind to interferon-stimulated response elements (ISREs) as a homodimer at loci where IRF4 is present at high concentration, whereas it remains unclear whether such a high local concentration of IRF4 is physiologically available *in vivo* (Ochiai et al., 2013). Wherefore, the homodimerization of IRF4, or its interactions with NFATc2 or STATs in CD8^+^ T cells, along with the impacts of BATF deficiency on the genomic bindings of those putative IRF4 partners should be explored in the future studies.

A previous study has shown that BATF and IRF4 are mutually dependent for their binding to DNA in TH17 cells (Ciofani et al., 2012). However, here we revealed that BATF bind to DNA and initiate chromatin reorganizations independent of IRF4, whereas interaction of IRF4 is crucial for following drastic alteration of the chromatin architecture and proliferation of effector CD8^+^ T cells. Collectively, these findings illustrated that BATF “pioneers” epigenomic remodeling to proceed the differentiation of cytolytic effector CD8^+^ T cells, as indicated in the differentiation of TH17 cells (Pham et al., 2019). However, the mechanisms by which BATF initiates the remodeling of poised chromatin remain unclear. Therefore, future works should examine epigenomic and transcriptomic landscapes at earlier time point (within hours after antigen encounter) to grasp the molecular events critical for fate decision driven by BATF.

We noted that BATF-bound loci were extensively co-localized by JunB, indicating the cooperative function of BATF and JunB. JunB plays crucial roles in T cell activation, differentiation of TH17 cells and Treg, through the interaction with various TFs such as NFAT proteins, Maf, and other AP-1 including BATF (Carr et al., 2017; Chinenov & Kerppola, 2001; Koizumi et al., 2018; Rengarajan et al., 2002). However, the role of JunB in CD8^+^ T cells has been ill-documented. Since the binding of JunB were considerably detected in the absence of BATF in our study, JunB may regulate CD8^+^ T cell differentiation in both BATF-dependent and independent manners. Thus, the involvement of JunB in CD8^+^ T cell differentiation, including the impact of JunB deficiency on the BATF binding should be addressed in future works.

Interestingly, the binding of JunD exhibited a substantial increase in *Batf*^-/-^ effector cells. Despite BATF and JunD have been shown to cooperatively function in CD4^+^ T cells (Bao et al., 2016; Kuwahara et al., 2016; Li et al., 2012), the role of JunD in CD8^+^ T cells remains poorly understood. A previous study indicated that the expression of JunD plays a critical role in transduction of IL-7 signaling (Ruppert et al., 2012). Given that BATF-deficient CD8^+^ T cells preferentially differentiate into IL-7R-expressing MPEC, JunD potentially contributes to the maintenance of MPEC-like effector CD8^+^ T cells while hampering the differentiation of cytolytic effector cells in the absence of BATF. Additionally, JunD promotes the expression of Nr4a1, a nuclear receptor that suppresses the proliferation and function of effector CD8^+^ T cells through the repression of IRF4 (X. Liu et al., 2019; Nowyhed et al., 2015; Stocco et al., 2002). Furthermore, it has been shown that JunD opposes JunB in human adenocarcinoma cell line (Selvaraj et al., 2015). These findings indicate that BATF acts as a negative regulator of JunD to reinforce the differentiation of cytolytic effector CD8^+^ T cells, as primarily suggested (Aronheim et al., 1997; Echlin et al., 2000). The role of JunD and the molecular mechanisms by which BATF and JunD reciprocally control their bindings in CD8^+^ T cell differentiation therefore emerges as an interesting issue for future studies. Of note, our data contradicted a previous report indicating a significant dependence of JunD binding on BATF (Tsao et al., 2022). There are two critical differences in experimental settings: firstly, Tsao et al. examined TF bindings in *in vitro*-activated CD8^+^ T cells, while we analyzed effector cells generated *in vivo*. Secondly, the previous study profiled the distribution of TFs by chromatin immunoprecipitation with sequence (ChIP-seq), which requires cross-linking of TFs to DNA, whereas we employed CUT&RUN, allowing for the assessment of TF-DNA interactions under native conditions. These observations underscore the importance of analyzing native samples generated *in vivo* to investigate the physiological roles of TFs. The mechanisms how BATF and IRF4 differentially regulate chromatin accessibilities remain elusive. A previous study revealed that the IRF4 complex harbored several epigenetic modulators, such as histone acetyltransferase and the nucleosome remodeling and deacetylase (NuRD) complex, in B cells (Ochiai et al., 2018). However, the constituents of the BATF-IRF4 complex in CD8^+^ T cells remain unexplored in detail, and the impact of IRF4 deficiency on the compositions of BATF complex have yet to be examined. Thus, investigation of BATF complex components using BATF-deficient CD8^+^ T cells transduced with wild-type BATF or H55Q mutant would provide novel insights into the effector differentiation programs initiated by BATF and IRF4. Remarkably, we found a profound decrease in the expression of Ezh2, the subunit of the Polycomb Repressive Complex 2 (PRC2) which catalyzes the trimethylation of histone 3 lysine 27 and thereby precluding gene expression, in *Batf*^-/-^ effector P14 cells. Moreover, the lack of BATF drastically hampered chromatin accessibility and the binding of TFs at the enhancer region of *Ezh2* gene. Several studies have shown that Ezh2 is required for the differentiation of cytolytic effector CD8^+^ T cells, through the epigenetic silencing of MPEC-associated genes (Gray et al., 2017; Kakaradov et al., 2017; Stairiker et al., 2020). Of note, Ezh2-deficient CD8^+^ T cells showed an intact proliferation kinetics at initial phase (∼48 h) of viral infection, but failed in subsequent proliferative burst, reflecting the trend observed in *Batf*^-/-^ effector cells (Kakaradov et al., 2017). Additionally, feedback regulation between IRF4 and Ezh2 was suggested in B cells (Ochiai et al., 2018), which may correspond to the previously defined model of the transcriptional circuit underlying effector differentiation of CD8^+^ T cells (Kurachi et al., 2014). Therefore, future studies are warranted to investigate the role of Ezh2 in BATF-dependent regulation of CD8^+^ T cell differentiation.

In the present study, we adoptively transferred an excess number of P14 cells for genomic analyses to obtain sufficient cells at day 3 post-infection. Although the data of both WT and *Batf*^-/-^ cells largely corresponded to the phenotypes of each population, we should be aware that these cells potentially underwent atypical differentiation kinetics. Therefore, future studies should analyze epigenomic and transcriptomic statuses using cells transferred at physiological numbers using single-cell-based RNA-seq, ATAC-seq, and CUT&Tag assays.

Taken together, we shed light on the mechanisms by which BATF regulates the differentiation of cytolytic effector CD8^+^ T cells in detail. BATF also contributes to the development and the function of exhausted T cells during chronic viral infection (Y. Chen et al., 2021; Quigley et al., 2010); however, its role in exhausted T cells has been controversial, since both overexpression and depletion of BATF countered exhaustion and enhanced cytotoxicity of tumor specific CD8^+^ T cells (Seo et al., 2021; X. Zhang et al., 2022). Therefore, future studies should investigate BATF-centric epigenomic and transcriptomic regulatory mechanisms at different stages of differentiation to comprehensively understand the role of BATF in CD8^+^ T cell differentiation.

## Methods

### Mice

WT C57BL/6 mice (CD45.1^+^) and Tg(Cd4-cre)1Cwi/BfluJ mice (CD4^Cre^, CD45.2^+^) were purchased from The Jackson Laboratory. B6.129S-*Batf^tm1.1Kmm^*/J mice (*Batf*^-/-^, CD45.2^+^) and B6.129S1-*Irf4^tm1Rdf^*/J mice (*Irf4*^fl/fl^, CD45.2^+^) were purchased from Charles River Labs Japan Inc. Each mouse was crossed with P14 mice transgenic for a TCR specific to the H-2D^b^ GP33-41 epitope of LCMV (CD45.2^+^, kindly provided from Dr. Ruka Setoguchi, RIKEN RCAI, Japan) to generate WT P14 (CD45.1^+^CD45.2^+^ or CD45.1^+^), CD4^Cre^ P14 (CD45.2^+^), *Batf*^-/-^ P14 (CD45.2^+^), and *Irf4*^fl/fl^ P14 (CD45.2^+^).

CD4^Cre^ P14 (CD45.2^+^), *Batf*^-/-^ P14 (CD45.2^+^), and *Irf4*^fl/fl^ P14 (CD45.2^+^) mice were crossed with WT P14 mice (CD45.1^+^) to generate CD4^Cre^ P14 (CD45.1^+^), *Batf*^-/-^ P14 (CD45.1^+^), and *Irf4*^fl/fl^ P14 (CD45.1^+^), respectively. CD4^Cre^ P14 (CD45.1^+^) and *Irf4*^fl/fl^ P14 (CD45.1^+^) mice were further crossed to generate IRF4 cKO mice (CD45.1^+^). WT C57BL/6 mice (CD45.2^+^) were purchased from Sankyo Labo Service Corporation, Inc. Male mice were used for experiments at between 6 and 10 weeks of ages. All animal experiments were approved by the committee on Animal Experimentation of Kanazawa University.

### LCMV infection

LCMV Armstrong strain was produced and titrated as previously described (Kurachi et al., 2014). Mice were infected with LCMV by intraperitoneal injection (2 × 10^5^ plaque-forming units).

### Adoptive transfer

For adoptive transfer of naive cells, CD8^+^ T cells were isolated from spleens of naive WT or *Batf*^-/-^ P14 mice using EasySep CD8 T cell isolation kit (STMCELL Technologies). The appropriate number of a mixture of cells (1:1 ratio; 1× 10^4^ to 1× 10^6^ each cell) or single P14 cells was intravenously transferred into nonirradiated naive recipient mice.

### Retroviral transduction

*Batf*, *Batf* H55Q mutant, *Batf* S43A mutant, *Batf* S43D mutant, *Batf* S43A/H55Q mutant cDNAs were cloned into a MSCV-IRES-VEX plasmid (kindly provided by Dr. Warren S. Pear, university of Pennsylvania) with Gateway Cloning Technology (Thermo Fisher Scientific). To produce retroviral supernatants, each plasmid was transfected into 293T human embryonic kidney cells in combination with pCL-Eco plasmid using jetOPTIMUS DNA Transfection Reagent (Polyplus) according to the manufacturer’s instructions. For the transduction of retrovirus, naive WT or *Batf*^-/-^ P14 cells were isolated from spleens as described above or using BD IMag^TM^ Mouse CD8 T Lymphocyte Enrichment Set (BD Biosciences), and stimulated for 24 h with plate-bound anti-CD3 (5 μg/ml; 2C11; Biolegend) and anti-CD28 (2 μg/ml; 37.51; Biolegend) in complete RPMI-1640 (RPMI-1640 medium supplemented with 10% fetal bovine serum (FBS), 20 mM HEPES, 50 μM 2-mercaptoethanol, 100 U/ml penicillin, 100 μg/ml streptomycin, 1 mM sodium pyruvate, 100 μM nonessential amino acids) containing 100 U/ml recombinant human IL-2 (Peprotech). P14 cells were then transduced with retroviral supernatant containing polybrene (8 μg/ml; Sigma-Aldrich) by spin infection (2,000*g* for 1 h at 30°C). Retroviral supernatants were transduced into P14 cells as previously described (Kurachi et al., 2017). After 4 h incubation, retrovirus-transduced P14 cells were adoptively transferred into recipient mice that had been infected with LCMV 1 d before, or cultured in complete RPMI-1640.

### Antibodies used in this study

The following antibodies were used for flow cytometry and cell sorting. From eBioscience/ThermoFisher Scientific: anti-CD8a-PE (53-6.7), dilution 1:100 or 1:200; anti-CD8a-NY700 (53-6.7), 1:200; anti-CD8a-APC (53-6.7), 1:200 or 1:300; anti-CD122-PerCP-EF710 (TM-brta1), 1:200; anti-CD127-BUV737 (A7R34), 1:200; anti-PD-1-FITC (RMP1-30), 1:200; anti-TIM3-SB702 (RMT3-23), 1:200; anti-CD45.1-BUV805 (A20), 1:100 or 1:200; anti-CD45.2-APC-eF780 (104), 1:200; anti-CD45.2-BUV395 (104), 1:100; anti-CD25-PE-Cy5 (PC61.5), 1:200; anti-CD44-FITC (IM7), 1:100; anti-CD49d-SB436 (R1-2), 1:200; anti-CD62L-BUV661 (MEL-14), 1:200; anti-IRF4-PerCP-eF710 (3E4), 1:100; anti-KLRG1-APC (2F1), 1:200; anti-KLRG1-PE-Cy7 (2F1), 1:200; anti-T-bet-PE-Cy7 (eBio4B10), 1:200; anti-TCR Va2-SB436 (B20.1), 1:100 or 1:200. From BD Biosciences: anti-CD45.1-BUV395 (A20), 1:200; anti-Ezh2-PE (11/EZH2), 1:100; anti-TCF-1/TCF7-AF488 (S33-966), 1:100. From Cell Signaling Technology: anti-BATF-AF647 (D7C5), 1:100; anti-BATF-PE (D7C5), 1:100; anti-TCF-1/TCF7-PE (C63D9), 1:100. From Biolegend: anti-CD45.1-Pacific Blue (A20), 1:200; anti-CD16/32 (93), 1:400.

The following antibodies were used for CUT&RUN assay. From Cell Signaling Technologies: anti-T-bet/TBX21 (E4I2K), 1:50; anti-JunB (C37F9), 1:50; anti-JunD (D17G2), 1:50; anti-FoxO1 (C29H4), 1:50. From Brookwood Biomedical: anti-BATF (PAB4003; lot 005), 0.5 μg/assay. From Santa Cruz: anti-IRF4 (E-7), 0.5 μg/assay. From Invitrogen: Rabbit IgG isotype control (cat# 02-6102), 0.5 μg/assay.

The following antibodies were used for EMSA. From Cell Signaling Technologies: anti-BATF (D7C5), 0.5 μl/assay; anti-IRF4 (D9P5H), 0.5 μl/assay.

### Flow cytometry and cell sorting

For flowcytometry, single-cell suspension from splenocytes were prepared by homogenizing spleens using a 70-μm cell strainer and treated with ACK lysis buffer to remove red blood cells. Cells were stained with Zombie Red Fixable Viability Kit (Biolegend, 1:2,000), Fixable Viability Dye eFluor 780 (eBioscience, 1:2,000), or Fixable Viability Stain 440 UV (BD Biosciences, 1:2,000) for 30 min at 4°C to exclude dead cells. Cells were then incubated with anti-CD16/32 antibodies for 15 min at 25°C to block Fc receptors, followed by surface staining with indicated antibodies for 30 min at 4°C. For the detection of TFs, surface-stained cells were fixed and permeabilized using eBioscience Foxp3/Transcription Factor Staining Buffer Set (Invitrogen) and stained with TF antibodies for 30 min at 4°C. Flow cytometry data were acquired on a BD FACSCanto II or BD FACSymphony A5 and analyzed with FlowJo v10 software (BD Biosciences).

For cell sorting, mononuclear cells were enriched from total splenocytes using Histopaque-1083 (Sigma-Aldrich) in accordance with the manufacturer’s instructions. Cells were incubated with Zombie Red Fixable Viability Kit for 30 min at 4°C and stained with indicated antibodies for 30 min at 4°C. P14 cells were sorted based on CD8, CD45.1, CD45.2, TCR-Va2, CD25, CD44, and GFP or violet-excited fluorescent protein (VEX) for IRF4 cKO cells or RV-transduced cells, respectively. Cell sorting was carried out on a BD FACSAria Fusion (BD Biosciences) in RPMI-1640 containing 50% FBS.

### Electrophoretic Mobility Shift Assay (EMSA)

Naive *Batf*^-/-^ P14 cells were isolated and transduced with empty RV, RV overexpressing *Batf*, or *Batf* H55Q mutant as described above. At day 3 after RV transduction, RV-transduced P14 cells were sorted based on the fluorescence of VEX reporter, and nuclear extracts were prepared using NE-PER™ Nuclear and Cytoplasmic Extraction Reagents (Thermo Fisher Scientific) in accordance with manufacturer’s instructions. Protein concentrations in each extract were quantified using Takara BCA Protein Assay Kit (Takara Bio). EMSA was performed using Gelshift™ Chemiluminescent EMSA (Active Motif) according to manufacturer’s protocols with modifications. Binding reactions contained 1 × binding buffer, 5 μg of nuclear extracts, 1 μg of poly d(I-C), 2.5% glycerol, 0.05% NP40, and 5 mM MgCl2. After 5 min incubation at 25°C, extracts were incubated with 0.5 μl of antibodies described above for 30 min at 4°C, followed by 30 min incubation with 10 fmol of biotinylated oligonucleotide containing AICE1 from third intron of mouse *Ctla4* (Tussiwand et al., 2012) (5′-CTT GCC TTA GAG GTT TCG GGA TGA CTA ATA CTG TA-3′/5′-TCA CGT ACA GTA TTA GTC ATC CCG AAA CCT CTA AGG-3′) at 25°C. Samples were electrophoresed on 4.5% polyacrylamide gels in 0.1 × TBE buffer and transferred onto Blotting-Nylon 66 membranes, type B, positive (Sigma-Aldrich). Oligonucleotide was then cross-linked to the membrane using UV transilluminator (Analytik Jena), and the membrane was incubated with Streptavidin HRP-conjugate for 15 min at 25°C. After exhaustive washing, the membrane was equilibrated in the substrate equilibration buffer and then reacted with the chemiluminescent reagent premixed with an equal volume of the reaction buffer. Signals were detected using the ChemiDoc MP Imaging System (Bio-rad).

### Sample preparation for RNA-seq

Freshly sorted cells were centrifuged and resuspended in RLT buffer (RNeasy Mini kit, QIAGEN) supplemented with 2-mercaptethanol and stored at -80°C. Total RNA was extracted using RNeasy Mini kit (QIAGEN) according to the manufacturer’s protocol. The quality of RNA was analyzed using a Bioanalyzer 2100 (Agilent), and libraries were prepared using NEBNext Ultra RNA LP Kit (New England Biolabs). Libraries were paired-end sequenced (150 bp + 150 bp) on a NovaSeq 6000 (Illumina).

### ATAC-seq library preparation and sequencing

ATAC-seq libraries were prepared as described (Buenrostro et al., 2015; Corces et al., 2017) with slight modifications. Approximately 2-5× 10^4^ cells were washed twice with cold PBS and resuspended in 50 μl of cold lysis buffer (10 mM Tris-HCl pH 7.5, 10 mM NaCl, 3 mM MgCl2, 0.1% Tween 20, 0.1% IGEPAL CA-630, and 0.01% digitonin). After 3 min incubation on ice, 1 ml of cold wash buffer (10 mM Tris-HCl pH7.5, 10 mM NaCl, 3 mM MgCl2, and 0.1% Tween 20) was added and lysates were centrifuged (500*g*, 10 min, 4°C). Supernatants were removed and isolated nuclei were resuspended in 50 μl of transposition mixture (25 μl of TD buffer, 2.5 μl of Tn5 transposase (Illumina), and 22.5 μl of nuclease-free water) and incubated for 30 min at 37°C. Transposed DNA fragments were then immediately purified using MinElute Reaction Cleanup Kit (QIAGEN). Sequence libraries were amplified using NEBNext High Fidelity 2× PCR Master Mix (New England Biolabs) and barcoded primers. The number of cycles for amplification was determined by a quantitative PCR side reaction. After amplification, libraries were size selected with AMPure XP (Beckman Coulter) beads using purification with bead/sample ratios of 1.8. The quality and quantity of libraries were determined by Tapestation 4150 (Agilent) and NEBNext Library Quant Kit for Illumina (New England Biolabs), respectively. Libraries were pooled equimolar and paired-end sequenced (67 bp + 155 bp) in the NovaSeq 6000 platform.

### Preparation of protein A/G-MNase protein

*E*. *coli* strain C3013 (New England Biolabs) harboring proein A/G-MNase-His (pAG-MNase) was cultured in 2YT medium containing 50 μg/ml kanamycin at 37°C. Expression of pAG-MNase was induced by supplementation of 0.1 mM isopropyl β-D-1-thiogalactopyranoside at OD600 0.6, followed by overnight culture at 18°C. Cells were harvested by centrifugation and resuspended in lysis buffer (50 mM HEPES pH8.0, 500 mM NaCl, 1 × CelLyticB (Sigma-Aldrich), 0.1 mM PMSF). The lysate was incubated at 37°C for 5 min and subjected to IMAC purification using TALON beads (Clontech). The eluate was concentrated by ultrafiltration using 10 kDa cut-off Vivaspin (Sartorius) and subjected to size exclusion chromatography using SperDex 200 increase column 10/300 (Sigma-Aldrich) with SEC buffer (20 mM HEPES pH7.5, 300 mM NaCl, 1 mM EDTA). After the concentration of the peak containing pAG-MNase by ultrafiltration using 10kDa cut-off Vivaspin (Sigma-Aldrich), the sample was mixed with equal volume of 100% glycerol and stored at -80°C until use.

### CUT&RUN library preparation and sequencing

CUT&RUN experiments were conducted as described (Meers et al., 2019), with modifications. Briefly, 1-4 × 10^5^ sorted cells were washed with cold phosphate-buffered saline (PBS) and nuclei were extracted by incubating cells in cold nuclear extraction buffer (20 mM HEPES-KOH pH 7.9, 10 mM KCl, 0.1% Triton X-100, 20% glycerol, 1 mM MnCl2, 0.5 mM spermidine, 1 × protease inhibitor cocktail (Roche)) on ice for 10 min. Nuclei were incubated with 10 μl of BioMagPlus Concanavalin A (Bang laboratories) at 4°C for 15 min. Tubes were placed on a magnetic stand and supernatants were removed. Primary antibodies in cold antibody buffer (20 mM HEPES pH 7.5, 150 mM NaCl, 0.5 mM spermidine, 0.1% bovine serum albumin (BSA), 1 × protease inhibitor cocktail, 2 mM EDTA, 0.01% digitonin) was added to the tubes and samples were incubated at 4°C overnight. The next day, cells were washed three times with cold Dig-wash buffer (20 mM HEPES pH 7.5, 150 mM NaCl, 0.5 mM spermidine, 0.05% BSA, 1× protease inhibitor cocktail, 0.01% digitonin), and pAG-MNase in cold Dig-wash buffer at 700 ng/ml was added to the tubes and rotated for 1 h at 4°C. Nuclei were washed three times with cold Dig-wash buffer and then rinsed with cold low-salt buffer (20 mM HEPES pH 7.5, 0.5 mM spermidine, 0.01% digitonin). pAG-MNase-mediated chromatin digestion was then initiated by incubating nuclei with cold MNase incubation buffer (3.5 mM HEPES pH 7.5, 10 mM CaCl2, 0.01% digitonin) for 30 min at 0°C. The tubes were placed on a chilled magnetic stand and the supernatants were removed. The reaction was terminated and the digested fragments were released into the solution by incubating nuclei in 1× stop buffer (170 mM NaCl, 20 mM EGTA, 0.01% digitonin, 50 μg/mL RNase A) at 37°C for 15 min. The tubes were placed on a magnetic stand and the supernatants were collected. CUT&RUN libraries were prepared using NEBNext Ultra II DNA Library Prep Kit for Illumina (New England Biolabs) according to the manufacturer’s protocols with modifications as described below. For dA-tailing, samples were incubated at 50°C to avoid melting of small DNA fragments and the incubation time was extended to 1 h (N. Liu et al., 2018). After the adaptor ligation, samples were purified with AMPure XP beads (Beckman Coulter) with a bead/sample ratio of 1.75 to remove excess adaptors and libraries were amplified by PCR. The number of PCR cycles was determined by a quantitative PCR side reaction. After amplification, libraries were size selected with AMPure XP beads using two consecutive rounds of purification with bead/sample ratios of 1.2. The quality and quantity of libraries were determined by Tapestation 4150 (Agilent) and NEBNext Library Quant Kit for Illumina (New England Biolabs), respectively. Libraries were pooled equimolar and paired-end sequenced on a NovaSeq 6000 (67 bp + 155 bp) or a NextSeq 2000 (50 bp + 50bp) (Illumina).

### RNA-seq data processing and analysis

Sequence data were assessed using FastQC, and trimmed for adapter sequences and low-quality bases using fastp(S. Chen, 2023). Reads were aligned to the GRCm38/mm10 reference genome using STAR (Dobin et al., 2013) with default settings. Read counts and transcripts per million (TPM) were calculated using featureCounts and StringTie (Pertea et al., 2015), respectively. Differentially expressed genes (DEGs) were identified using DESeq2 v1.42.0, and genes were considered as differentially expressed if they have an FDR < 0.05 and fold change > 2.0 or <-2.0. GSEA was performed using the Broad Institute software (https://www.gsea-msigdb.org/gsea/index.jsp) or fgsea v1.28.0 for the ImmSigDB C7 and GO Biological Processes (C5) collections. Data were visualized using Prism9 (GraphPad Software) or R.

### ATAC-seq data processing and analysis

Sequence data were assessed using FastQC, and trimmed for adapter sequences and low-quality bases using fastp(S. Chen, 2023). Reads were aligned to the GRCm38/mm10 reference genome using Bowtie2 v2.3.5. To remove unmapped, unpaired, low-quality reads (MAPQ < 30), and mitochondrial reads, samtools was used. ENCODE blacklist regions were removed, and PCR duplicates were removed using Picard. BED file was created using bamtobed function of BedTools and ATAC shift was corrected as described (https://github.com/wherrylab/jogiles_ATAC). Peak calling was performed using MACS v2 using the narrowPeak setting with q value cut-off of 0.01, -f BEDPE. Peaks from all samples in each experiment were combined and merged using BedTools merge, and the numbers of reads in each peak was quantified using BedTools coverage. Genes proximal to peaks were annotated against the mm10 genome using ChIPseeker v1.38.0. Differences in accessibilities of peaks were analyzed using DESeq2 v1.42.0 normalization. Peaks were considered as differentially accessible region if they have a false-discovery rate (FDR) < 0.05 and fold change > 2.0 or <-2.0. Tracks were visualized using Integrative Genomic Viewer (v v2.12.3). Gene ontology term and gene set enrichment analyses were performed using rGREAT package v2.4.0 (Gu & Hübschmann, 2023), for the ImmSigDB C7 collection. Signal heatmaps were created using deepTools v3.5.4 with computeMatrix function using differentially accessible regions or shared accessible regions as reference files, with following settings: reference-point -referencePoint center -a 1000 -b 1000 --missingDataAsZero --SkipZero. Overlaps in peaks between samples were determined using mergePeaks function of HOMER or BedTools intersect. Motif enrichment analysis was performed using the findMotifsGenome.pl function of HOMER (Heinz et al., 2010). Data were visualized using Prism9 (GraphPad Software) or R.

### CUT&RUN data processing and analysis

Sequence data were assessed using FastQC, and trimmed for adapter sequences and low-quality bases using fastp(S. Chen, 2023). Reads were then aligned to the GRCm38/mm10 reference genome using Bowtie2 v2.3.5. To remove unmapped, unpaired, low-quality reads (MAPQ < 30), and mitochondrial reads, samtools was used.

ENCODE blacklist regions were removed, and PCR duplicates were removed using Picard. Filtrated bam files biological replicates were merged, and following procedures were performed both for each biological replicate file and merged files. BED file was created using bamtobed function of BedTools. Data were normalized by mapping the reads to *E*. *coli* DNA, the carry-over from purification of pAG-MNase (Meers et al., 2019) and normalized bigwig files were created using deepTools bamCoverage. The bigwig files from each biological replicate were analyzed using multiBigwigSummary function of deepTools to confirm the reproducibility of data. Peak calling was performed using MACS v2 using the narrowPeak setting for TFs with q value cut-off of 0.01, -f BEDPE. For the background, the bed file of IgG CUT&RUN data was used to identify target-specific signals unless otherwise indicated. Peak overlaps between samples were determined using mergePeaks function of HOMER.

Peaks from all samples in each experiment were combined and merged using BedTools merge, and the numbers of reads in each peak was quantified using BedTools coverage. Signal density heatmap was generated using deepTools v3.5.4 as described above using merged peaks, or differentially accessible regions or shared accessible regions determined by ATAC-seq as reference files. Genes proximal to peaks were annotated against the mm10 genome using ChIPseeker v1.38.0. Differences in bindings between samples were analyzed using DiffBind v3.12.0. Peaks were considered as differentially bound region if they have an FDR < 0.05. Tracks were visualized using Integrative Genomic Viewer (v2.12.3). Motif enrichment was analyzed using the findMotifsGenome.pl function of HOMER. Data were visualized using Prism9 (GraphPad Software) or R.

### Statistical analysis

For comparison of data between two groups, an unpaired, two-tailed Student’s *t*-test was performed. For multiple comparison of data, Kruskal-Wallis test was carried out. Analyses were performed using Prism9. Information for statistical analyses of genomic data is provided in the corresponding methods section.

## Acknowledgements

This work was supported in part by Grants-in-Aid for Scientific Research under Grants (C: 19K07476 and B: 23K24109) and Grant-in-Aid for Early-Career Scientists (23K14538) from the Japan Society for the Promotion of Science, AMED under grant number JP24fk0310509, and Kanazawa University SAKIGAKE project. This work was the result of using research equipment shared in MEXT Project for promoting public utilization of advanced research infrastructure (Program for supporting construction of core facilities) Grant Number JPMXS04403000XX.

## Author contributions

SF, YT, AH, RS, and TT carried out the experiments. JK, MK, and YM provided key resources. SF and YT analyzed and visualized data. YM, EJW, and MK provided intellectual input. MK conceived of the project. SF, YT, and MK drafted the manuscript. All authors have reviewed and approved the final manuscript.

