## Supplementary materials for "The transcription factor BATF pioneers the differentiation program of cytolytic effector CD8^+^ T cells through the direct interaction with IRF4"

Supplementary figure 1

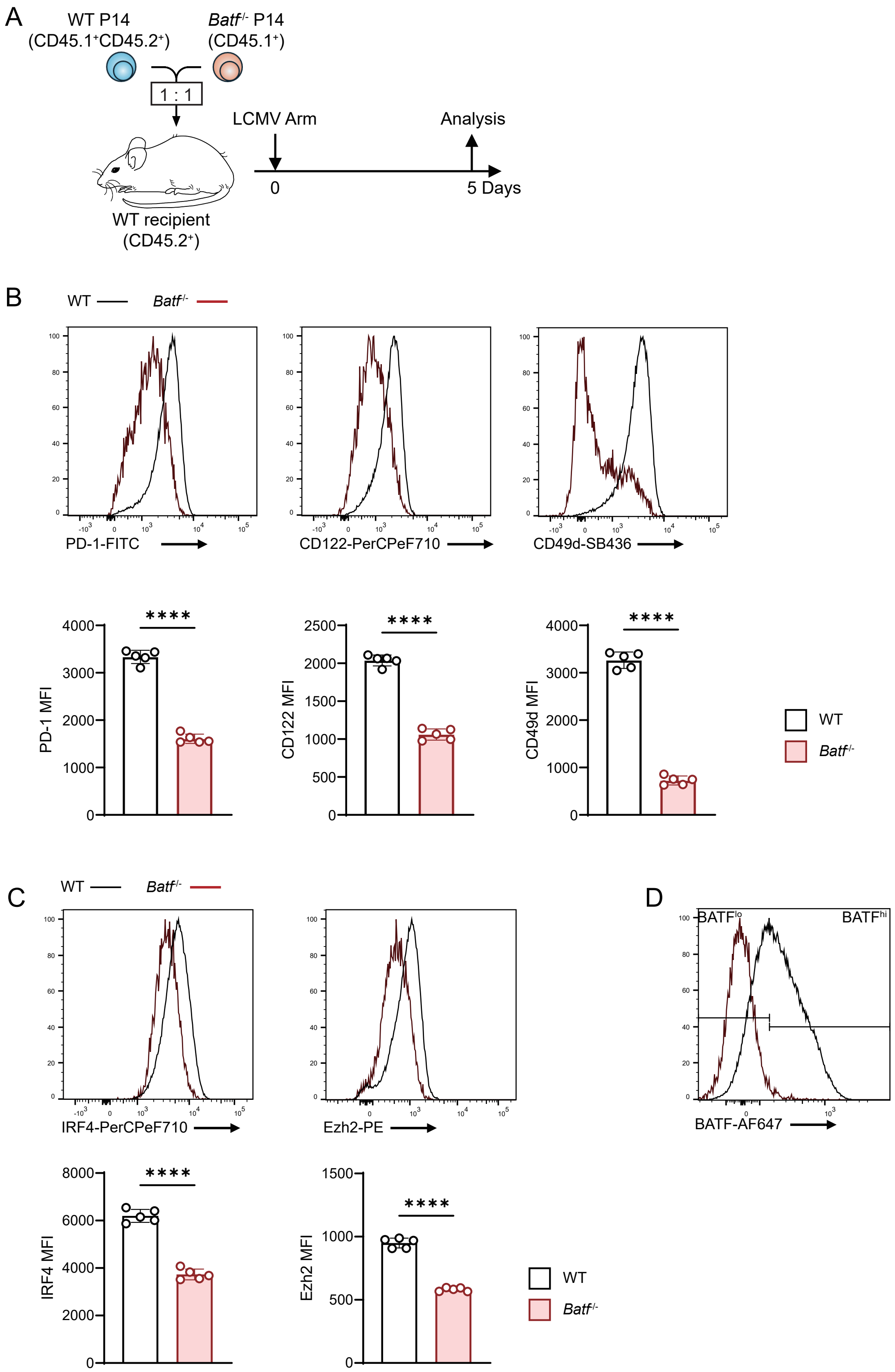

**Supplementary figure 1. BATF-deficient CD8+ T cells exhibited decreased expression of signatures associated with SLEC phenotype.**

(A) Experimental approaches. WT recipient mice were adoptively transferred with congenically distinct WT and *Batf* $^{-/-}$ P14 cells mixed at 1:1 ratio ($1 \times 10^4$ each cell), followed by infection of the recipient mice with LCMV Arm, analyzed at day 5 post-infection. (B, C) Flow cytometric analysis of splenocytes of the recipient mice in A. Plots are gated on each P14 cell. (B) Expression of PD-1, CD122, and CD49d, and (C) expression of IRF4 and Ezh2. Bar plots indicate the mean fluorescence intensity (MFI) of indicated markers in each P14 cell. (D) A gating strategy to classify WT P14 cells into BATF$^{hi}$ and BATF$^{lo}$ populations analyzed in Fig. 1F. ****$p < 0.0001$ (unpaired Student' s *t*-test). Data are representative of two independent experiments with four to five mice in each experiment.

### Supplementary figure 2

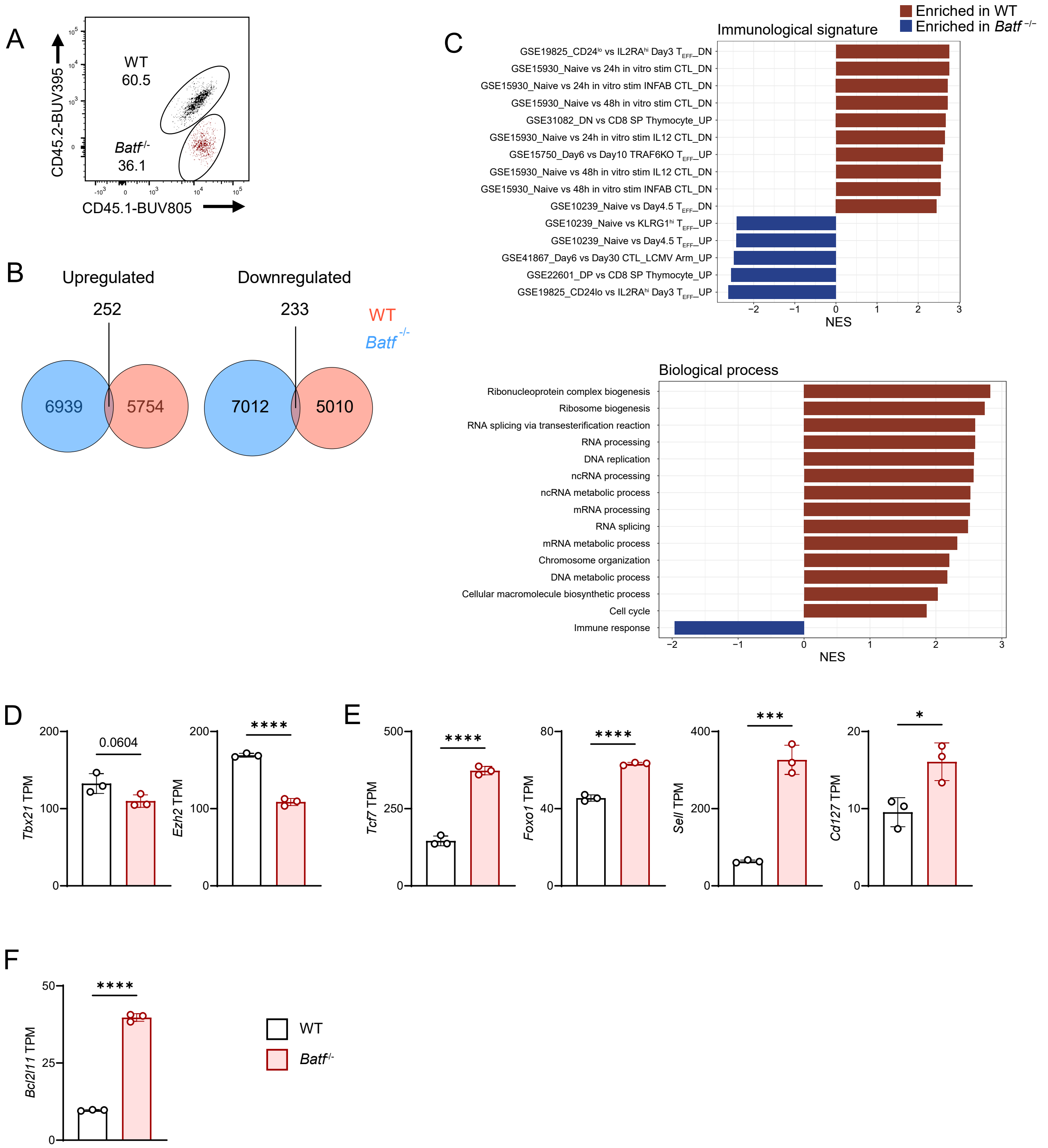

**Supplementary figure 2. Lack of BATF disrupt the transcriptional reprogramming during early effector differentiation of CD8$^+$ T cells.**

WT recipient mice were adoptively transferred with congenically distinct WT and *Batf* $^{-/-}$ P14 cells mixed at 1:1 ratio (1 × $10^6$ each cell), followed by infection of the recipient mice with LCMV Arm. WT and *Batf* $^{-/-}$ P14 cells collected from spleen of the recipient mice at day 3 post-infection ( “effector” ) and naive WT and *Batf* $^{-/-}$ P14 cells were subjected to RNA-seq (effector, *n* = 3; naive, *n* = 2). (A) Flow cytometric analysis of splenocytes obtained from the recipient mice at day 3 post-infection. Plots are gated on P14 cells. (B) Venn diagrams represent the numbers of genes upregulated (left) and downregulated (right) during transition from naive to effector in WT or *Batf* $^{-/-}$ P14 cells. (C) GSEA between WT and *Batf* $^{-/-}$ effector P14 cells of the ImmSigDB C7 (top) and GO Biological Processes C5 (bottom). NES, normalized enrichment score. (D-F) Expression of (D) SLEC-related gens *Tbx21*, *Ezh2*, (E) MPEC-related genes *Tcf7*, *Foxo1*, *Sell*, *CD127*, and (F) pro-apoptotic gene *Bcl2l11* in effector P14 cells. *$p < 0.05$, **$p < 0.01$, ***$p < 0.001$, and ****$p < 0.0001$ (unpaired Student’ s *t*-test).

Supplementary figure 3

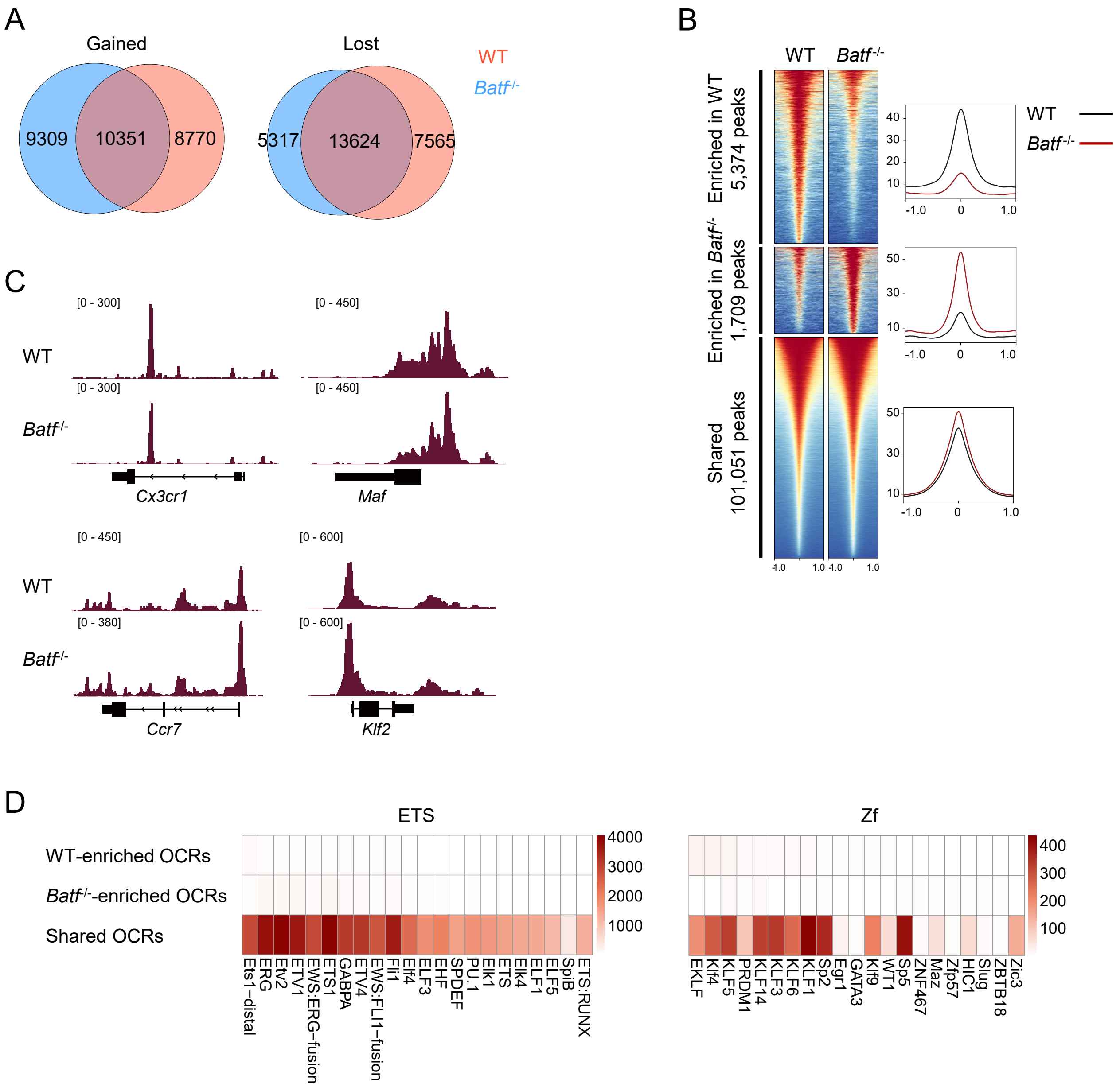

**Supplementary figure 3. BATF-deficient CD8$^+$ T cells manifest dysregulated chromatin remodeling in response to acute viral infection.**

WT recipient mice were adoptively transferred with congenically distinct WT and *Batf* $^{-/-}$ P14 cells mixed at 1:1 ratio (1 × $10^6$ each cell), followed by infection of the recipient mice with LCMV Arm. WT and *Batf* $^{-/-}$ P14 cells collected from spleen of the recipient mice at day 3 post-infection ( “effector” ) and naive WT and *Batf* $^{-/-}$ P14 cells were subjected to ATAC-seq ($n$ = 2). (A) Venn diagrams represent the numbers of OCRs increased (left) and decreased (right) their accessibilities during transition from naive to effector in WT or *Batf* $^{-/-}$ P14 cells. (B) ATAC-seq signal coverages at OCRs differentially accessible or shared between WT and *Batf* $^{-/-}$ effector P14 cells. Horizontal axis represents distance from peak center (kb). (C) Representative ATAC-seq signal tracks for WT and *Batf* $^{-/-}$ effector P14 cells. (D) Enrichment of selected TF motifs in differentially accessible or shared OCRs between WT and *Batf* $^{-/-}$ effector P14 cells. Color bars represent enrichment scores (-log10 $p$-value).

Supplementary figure 4

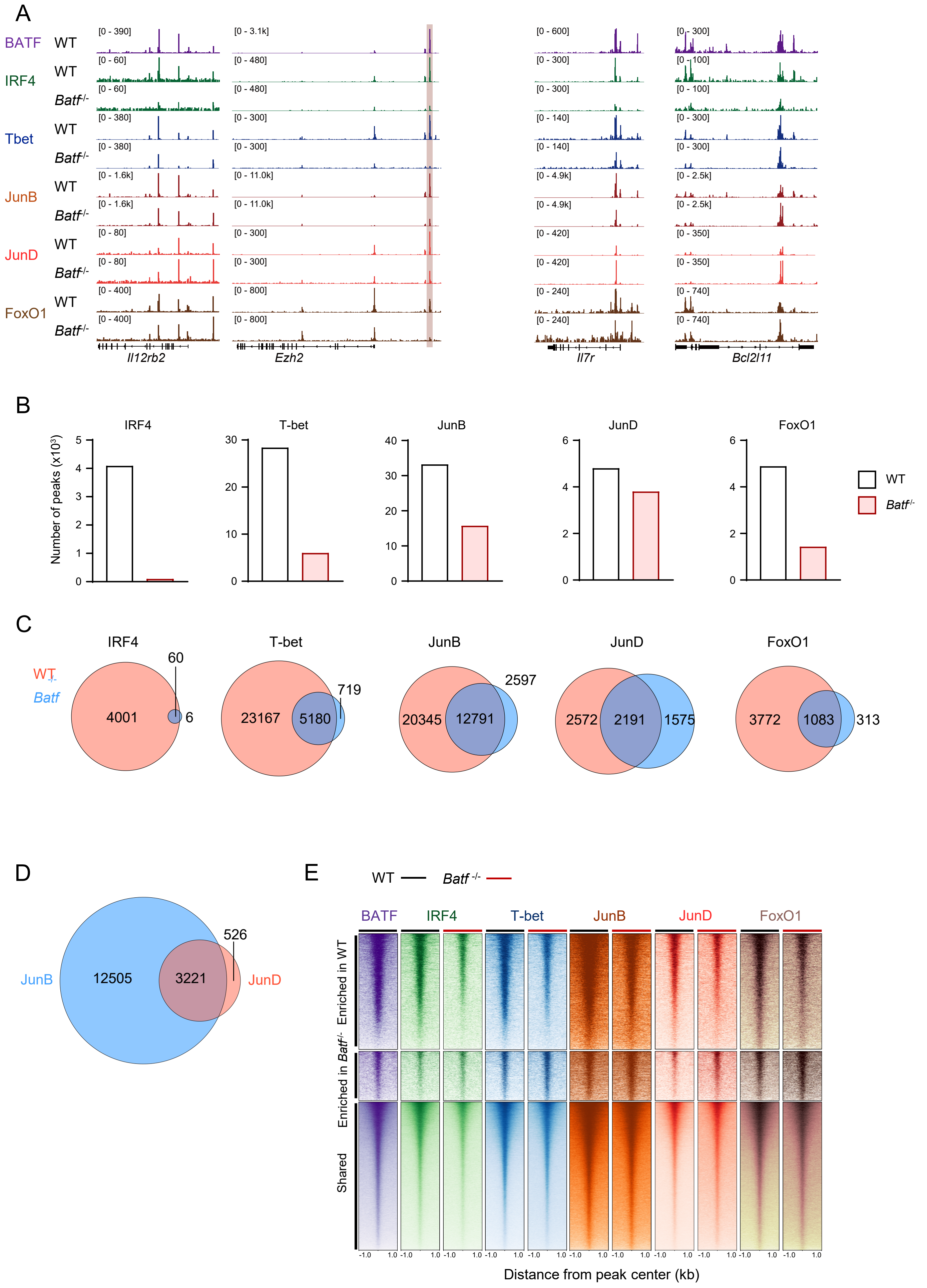

**Supplementary figure 4. BATF regulates the bindings of key transcription factors during effector CD8$^+$ T cell differentiation.**

WT recipient mice were adoptively transferred with congenically distinct WT and *Batf* $^{-/-}$ P14 cells mixed at 1:1 ratio (1 × $10^6$ each cell), followed by infection of the recipient mice with LCMV Arm. WT and *Batf* $^{-/-}$ effector P14 cells were collected from spleen of the recipient mice at day 3 post-infection and subjected to CUT&RUN (*n* = 2) analysis. Data from biological replicates were merged and analyzed. (A) Representative CUT&RUN signal tracks for WT and *Batf* $^{-/-}$ effector P14 cells. (B, C) The numbers of peaks detected in each CUT&RUN experiment visualized in (B) bar plots and (C) Venn diagrams. (D) Venn diagram showing loci bound by JunB and JunD in *Batf* $^{-/-}$ effector P14 cells. (E) Heatmap represents CUT&RUN signal coverages at OCRs differentially accessible or shared between WT and *Batf* $^{-/-}$ effector P14 cells in Fig. 2.

Supplementary figure 5

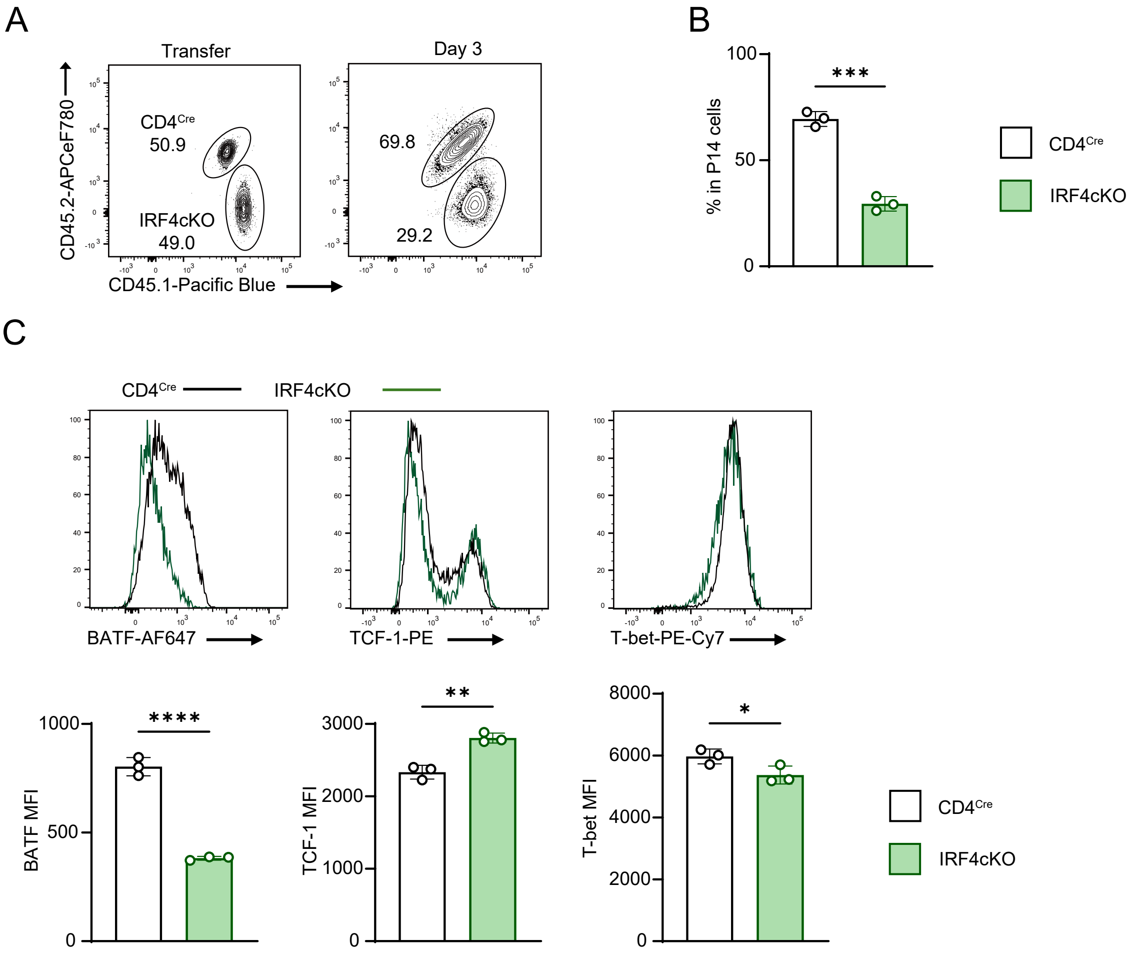

**Supplementary figure 5. Loss of IRF4 substantially impairs the differentiation of effector $CD8^+$ T cells.**

(A-C) Flow cytometric analysis of cells obtained from spleen of WT recipient mice adoptively transferred with congenically distinct $CD4^{Cre}$ and IRF4cKO P14 cells mixed at 1:1 ratio ($1 \times 10^6$ each cell), followed by infection of the recipient mice with LCMV Arm ($n = 3$). (A) Representative plots showing frequencies of $CD4^{Cre}$ and IRF4cKO P14 cell among total P14 cells analyzed at transfer (left) and day 3 post-infection (right). Plots are gated on P14 cells. (B) Frequencies of $CD4^{Cre}$ and IRF4cKO P14 cells at day 3 post-infection. (C) Expression of BATF, TCF-1, and T-bet in each effector P14 cell. Bar plots represents MFI of each TF. *$p < 0.05$, **$p < 0.01$, and ****$p < 0.0001$ (unpaired Student' s *t*-test).

### Supplementary figure 6

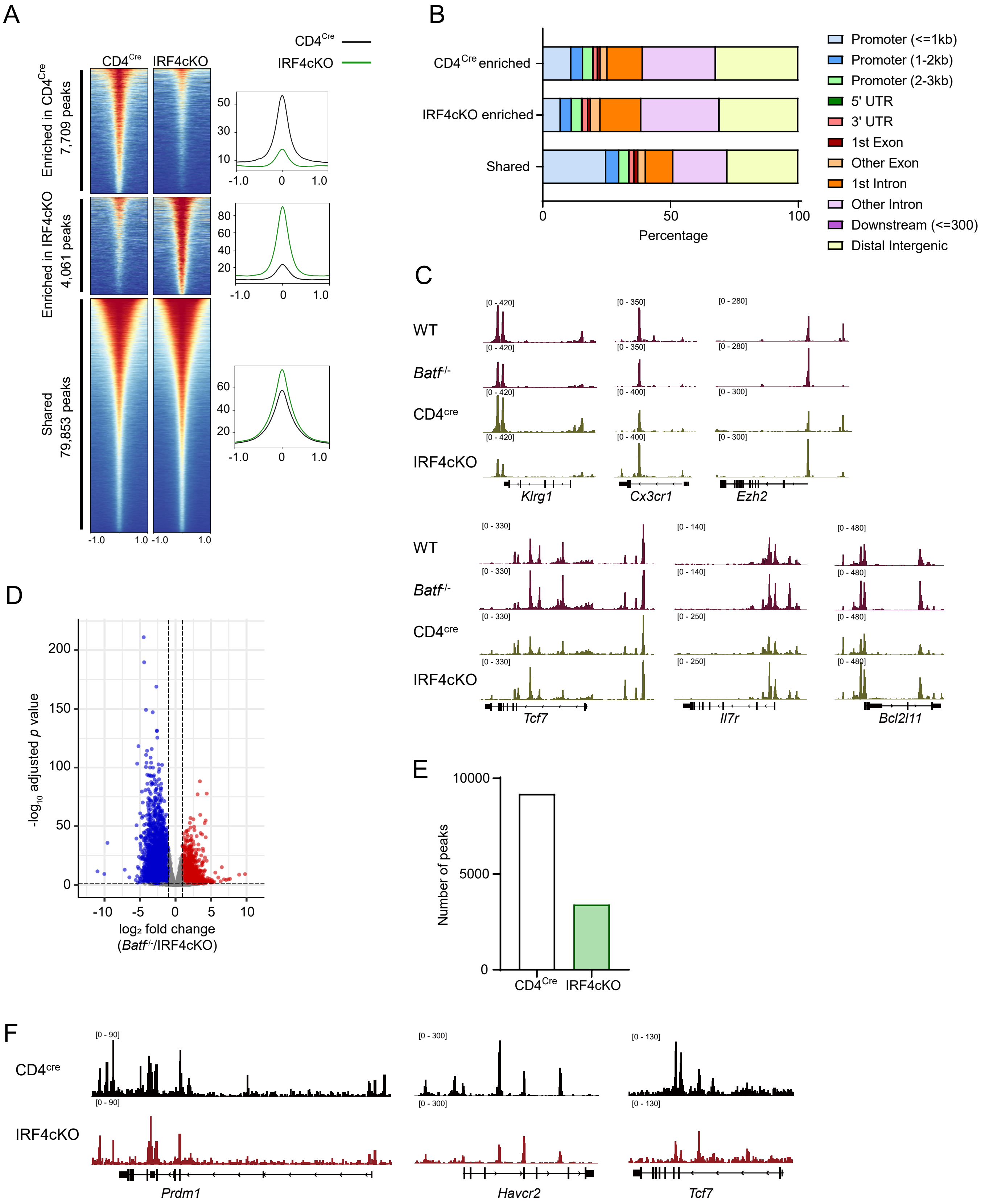

**Supplementary figure 6. Loss of IRF4 perturbs the chromatin accessibility and the binding of BATF in effector CD8$^+$ T cells.**

WT recipient mice were adoptively transferred with congenically distinct CD4$^{Cre}$ and IRF4cKO P14 cells mixed at 1:1 ratio (1 × $10^6$ each cell), followed by infection of the recipient mice with LCMV Arm. CD4$^{Cre}$ and IRF4cKO effector P14 cells were collected from spleen of the recipient mice at day 3 post-infection and subjected to ATAC-seq and BATF CUT&RUN analyses (*n* = 2). (A) ATAC-seq signal coverages at OCRs differentially accessible or shared between CD4$^{Cre}$ and IRF4cKO effector P14 cells. Horizontal axis represents distance from peak center (kb). (B) Peak annotation of OCRs differentially accessible or shared between CD4$^{Cre}$ and IRF4cKO effector P14 cells. (C) Representative ATAC-seq signal tracks for WT, *Batf* $^{-/-}$ (red), CD4$^{Cre}$, and IRF4cKO (yellow) effector P14 cells. (D) Volcano plot showing differences in chromatin accessibilities between *Batf* $^{-/-}$ and IRF4cKO effector P14 cells. (E) The numbers of BATF-bound peaks detected in a CUT&RUN experiment. (F) Representative BATF CUT&RUN signal tracks for CD4$^{Cre}$ and IRF4cKO effector P14 cells. (E, F) Data from biological replicates were merged and analyzed.

Supplementary figure 7

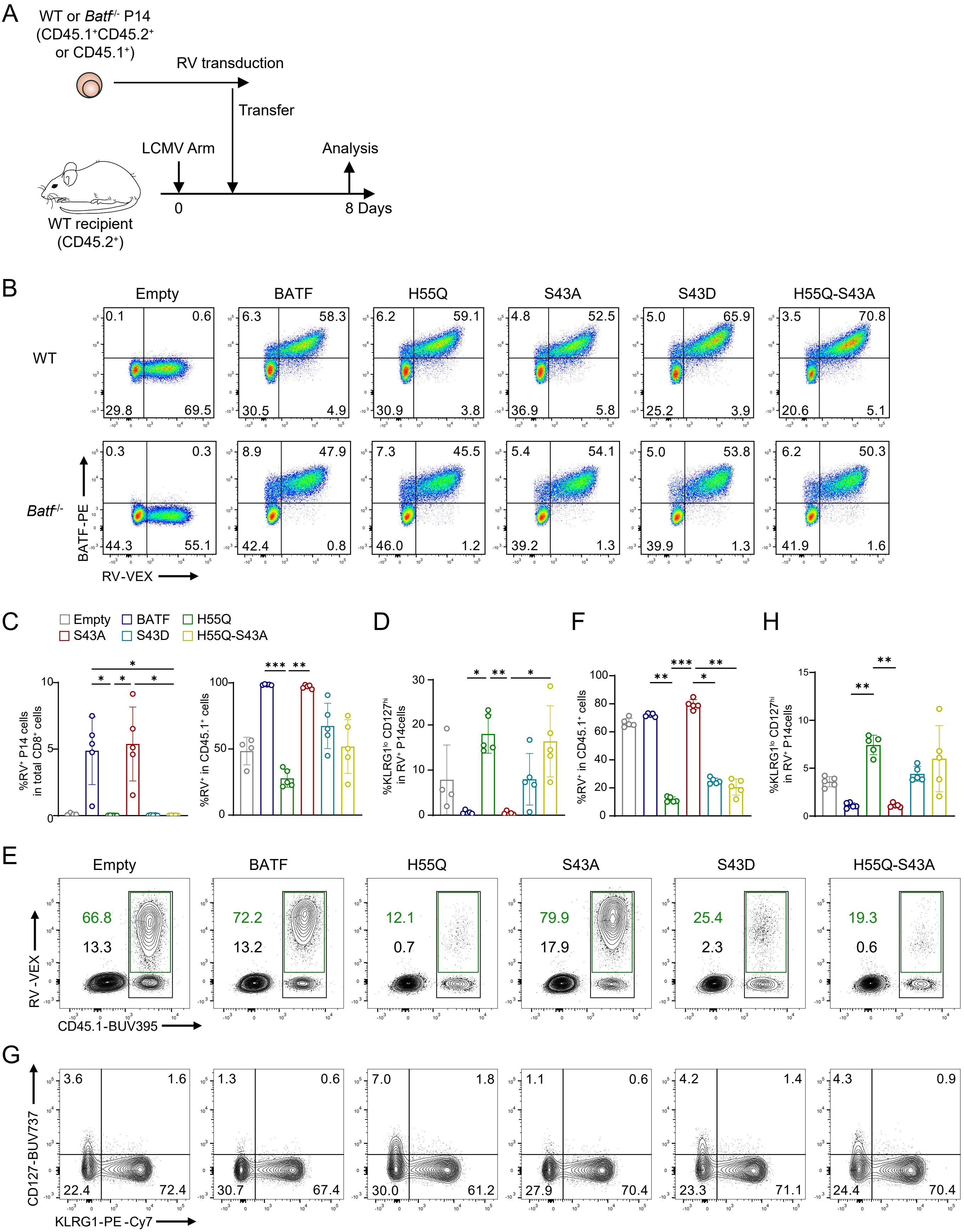

**Supplementary figure 7. Cooperation with IRF4 is crucial for BATF-mediated effector CD8$^+$ T cell differentiation.**

WT recipient mice were infected with LCMV Arm, and adoptively transferred with *Batf* $^{-/-}$ (C, D) or WT (E-H) P14 cells (1 × 10$^5$ each cell) transduced with empty retrovirus (RV) or RV overexpressing *Batf*, *Batf* H55Q, *Batf* S43A, *Batf* S43D, or *Batf* H55Q-S43A after 1 d stimulation in vitro with anti-CD3 and anti-CD28 antibodies. (A) Experimental approaches. (B-H) Flow cytometric analysis of splenocytes of the recipient mice in A, assessed at day 8 post-infection. (B) RV-transduced cells were cultured *in vitro* for 1 day after RV transduction, and efficiency of RV transduction and expression of BATF were analyzed. Plots are gated on P14 cells. (C) Percent WT P14 cells among total CD8$^+$ T cells (left) and percent RV-transduced violet-excited fluorochromes (VEX)$^+$ cells among total *Batf* $^{-/-}$ P14 cells (right). (D) Percent KLRG1$^{lo}$ CD127$^{hi}$ MPEC populations among total RV$^+$ *Batf* $^{-/-}$ P14 cells. (E) Representative plots showing the frequency of WT P14 cells and RV$^+$ cells among CD8$^+$ T cells. Plots are gated on total CD8$^+$ T cells and numbers indicate percent WT P14 cells among total CD8$^+$ T cells (black) and percent RV-transduced VEX$^+$ cells among total WT P14 cells (green). (F) Percent RV-transduced VEX$^+$ cells among total WT P14 cells. (G) Expression of KLRG1 and CD127 on RV-transduced WT P14 cells; plots are gated on VEX$^+$ P14 cells. (H) Percent KLRG1$^{lo}$ CD127$^{hi}$ MPEC populations among total RV$^+$ WT P14 cells. (B-H) Data are representative of two independent experiments with three to five mice in each experiment. (C, D, F, H) *$p < 0.05$, **$p < 0.01$, and ***$p < 0.001$ (Kruskal-Wallis test).

Supplementary figure 8

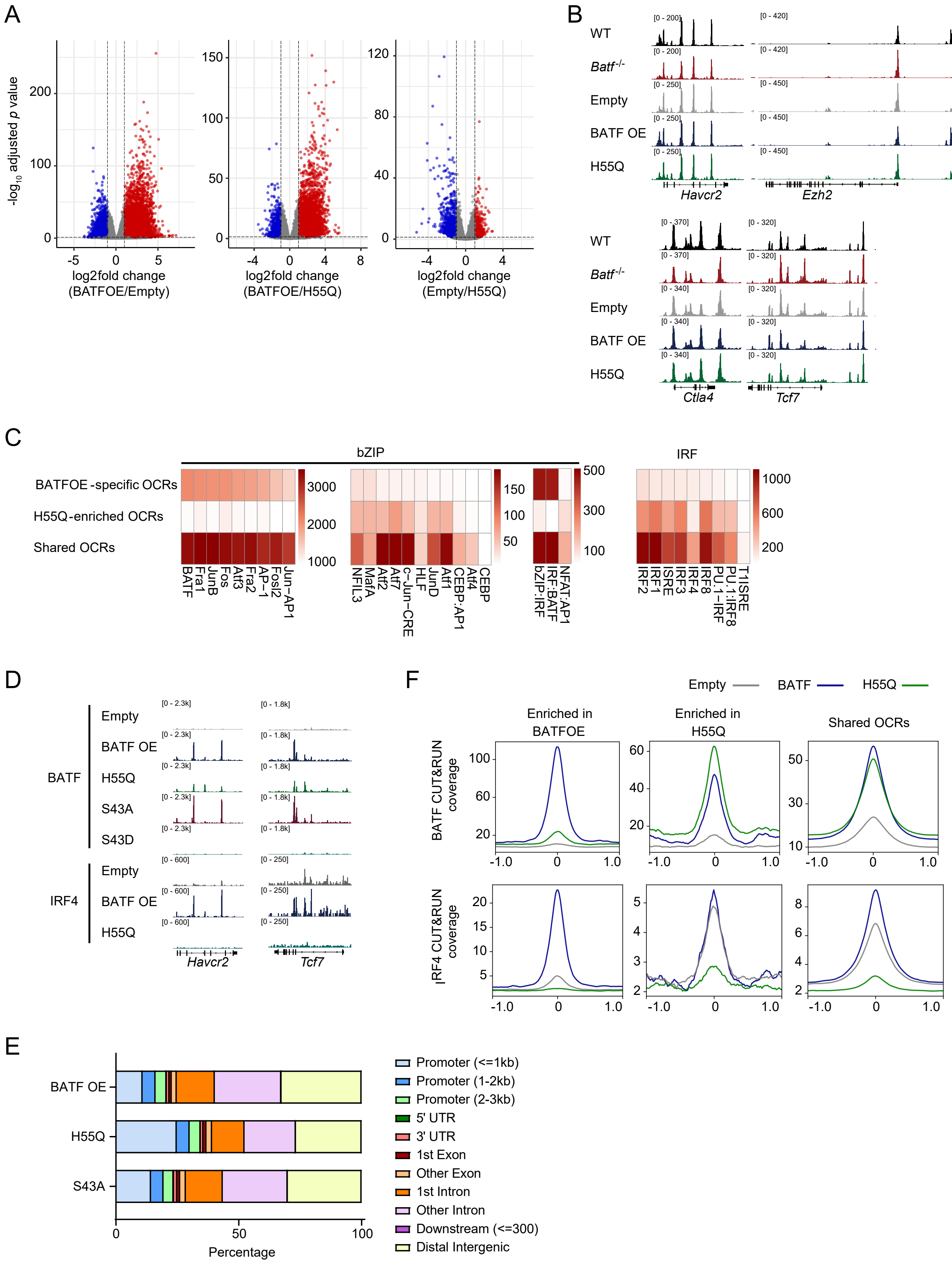

**Supplementary figure 8. BATF-IRF4 interaction is required for proper reprogramming of the chromatin landscape in CD8$^{+}$ T cells.**

(A, B) WT recipient mice were infected with LCMV Arm, and adoptively transferred with *Batf* $^{-/-}$ P14 cells (1 × 10$^{5}$ each cell) transduced with empty RV or RV overexpressing *Batf* or *Batf* H55Q after 1 d stimulation in vitro with anti-CD3 and anti-CD28 antibodies. RV$^{+}$ P14 cells were collected from spleen of the recipient mice at day 3 post-infection and subjected to ATAC-seq analysis (*n* = 2) (A) Volcano plots showing differences in chromatin accessibility between *Batf* RV- and empty RV-transduced cells (left), *Batf* RV- and H55Q RV-transduced cells (center), and empty RV- and H55Q RV-transduced cells (right). Horizontal axis represents fold change in chromatin accessibilities and vertical axis indicates statistical significance (log10 adjusted *p*-value). (B) Representative ATAC-seq signal tracks for WT, *Batf* $^{-/-}$ effector P14 cells (Fig. 2), and *Batf* $^{-/-}$ effector P14 cells transduced with indicated RV. (C) Enrichment of selected TF motifs in BATF-bound loci specifically observed or shared between Batf- and H55Q-expressing cells. Color bars represent enrichment scores (-log10 *p*-value). (D-F) *Batf* $^{-/-}$ P14 cells transduced with empty RV, RV overexpressing *Batf*, *Batf* H55Q, S43A, or S43D mutant were cultured *in vitro*, and subjected to CUT&RUN analysis at day 3 after RV transduction (*n* = 2). Data from biological replicates were merged and analyzed. (D) Representative BATF (upper) or IRF4 (lower) CUT&RUN signal tracks for *Batf* $^{-/-}$ P14 cells transduced with indicated RV. (E) Peak annotation of BATF-bound loci in cells transduced with RV overexpressing Batf, H55Q, or S43A. (F) Signal coverages of BATF (upper) and IRF4 (lower) CUT&RUN of *Batf* $^{-/-}$ P14 cells transduced with indicated RV at OCRs differentially accessible or shared between *Batf* RV- or H55Q RV-transduced effector P14 cells *in vivo*. Horizontal axis represents distance from peak center (kb).

**Supplementary Table 1. GSEA of the apoptotic processes between WT and *Batf*$^{-/-}$ effector P14 cells.**

| Pathway | padj | NES (WT vs *Batf*$^{-/-}$) |
|---|---|---|
| LEUKOCYTE_APOPTOTIC_PROCESS | 2.77E-03 | -1.75 |
| T_CELL_APOPTOTIC_PROCESS | 1.22E-02 | -1.72 |
| REGULATION_OF_LEUKOCYTE_APOPTOTIC_PROCESS | 2.88E-02 | -1.64 |
| LYMPHOCYTE_APOPTOTIC_PROCESS | 2.99E-02 | -1.63 |
| NEGATIVE_REGULATION_OF_LEUKOCYTE_APOPTOTIC_PROCESS | 3.30E-02 | -1.60 |
| NEGATIVE_REGULATION_OF_APOPTOTIC_SIGNALING_PATHWAY | 3.47E-02 | 1.40 |
| POSITIVE_REGULATION_OF_APOPTOTIC_SIGNALING_PATHWAY | 3.87E-02 | -1.49 |
| NEGATIVE_REGULATION_OF_INTRINSIC_APOPTOTIC_SIGNALING_PATHWAY | 4.41E-02 | 1.49 |

WT and *Batf*$^{-/-}$ effector P14 cells were subjected to RNA-seq as in Fig. 2. Enrichment of genes associated with apoptosis among GO Biological Processes (C5) were analyzed.
Gene sets of which adjusted *p*-value < 0.05 were listed. padj, adjusted *p*-value; NES, normalized enrichment score.

**Supplementary Table 2. Motif enrichment of BATF-bound loci.**

| Rank | Motifs | log10 *p*-value |
|---|---|---|
| BATF OE-specific peaks | | |
| 1 | BATF (GSE39756) | -2.606e+03 |
| 2 | Fos (GSE110950) | -2.505e+03 |
| 3 | JunB (GSE36099) | -2.492e+03 |
| 4 | Atf3 (GSE33912) | -2.438e+03 |
| 5 | Fra1 (GSE46166) | -2.397e+03 |
| ⋮ | | |
| 20 | IRF:BATF (GSE66899) | -4.338e+02 |
| H55Q-specific peaks | | |
| 1 | BATF (GSE39756) | -1.586e+02 |
| 2 | JunB (GSE36099) | -1.584e+02 |
| 3 | Atf3 (GSE33912) | -1.549e+02 |
| 4 | Fos (GSE110950) | -1.540e+02 |
| 5 | Fra1 (GSE46166) | -1.499e+02 |
| Shared peaks | | |
| 1 | JunB (GSE36099) | -2.445e+03 |
| 2 | BATF (GSE39756) | -2.413e+03 |
| 3 | Fos (GSE110950) | -2.355e+03 |
| 4 | Fra1 (GSE46166) | -2.336e+03 |
| 5 | Atf3 (GSE33912) | -2.314e+03 |
| ⋮ | | |
| 64 | IRF:BATF (GSE66899) | -4.853e+01 |

*Batf* $^{-/-}$ P14 cells transduced with retrovirus overexpressing *Batf* (BATF OE) or *Batf* H55Q mutant in Fig. 5. Enrichment of TF motifs in BATF-bound peaks specifically detected in BATF OE or H55Q samples and those shared between each cell.

**Supplementary Table 3. Motif enrichment of IRF4-bound loci.**

| Rank | Motifs | log10 *p*-value |
|---|---|---|
| BATF OE-specific peaks | | |
| 1 | BATF (GSE39756) | -1.805e+03 |
| 2 | JunB (GSE36099) | -1.704e+03 |
| 3 | Fos (GSE110950) | -1.616e+03 |
| 4 | Atf3 (GSE33912) | -1.600e+03 |
| 5 | AP-1 (GSE21512) | -1.567e+03 |
| ⋮ | | |
| 16 | IRF:BATF (GSE66899) | -3.533e+02 |
| Empty-specific peaks | | |
| 1 | Sp1 | -5.158e+01 |
| 2 | Klf1 (GSE136251) | -4.521e+01 |
| 3 | Klf3 (GSE44748) | -3.872e+01 |
| 4 | Klf5 (GSE49402) | -3.757e+01 |
| 5 | Klf6 (GSE64557) | -3.674e+01 |
| Shared peaks | | |
| 1 | BATF (GSE39756) | -1.647e+02 |
| 2 | JunB (GSE36099) | -1.549e+02 |
| 3 | Atf3 (GSE33912) | -1.419e+02 |
| 4 | AP-1 (GSE21512) | -1.371e+02 |
| 5 | Fos (GSE110950) | -1.361e+02 |
| ⋮ | | |
| 14 | IRF:BATF (GSE66899) | -7.484e+01 |

*Batf* $^{-/-}$ P14 cells transduced with retrovirus (RV) overexpressing *Batf* (BATF OE) empty RV as in Fig. 5. Enrichment of TF motifs in IRF4-bound peaks specifically detected in BATF OE or empty samples and those shared between each cell.
